# Neural subspace reorganization reflects value-based decision making

**DOI:** 10.64898/2026.02.02.703171

**Authors:** Huidi Li, Nikolaos Chrysanthidis, Scott L Brincat, Jonas Rose, Earl K Miller

## Abstract

Value-based decision making requires maintaining and comparing multiple option representations associated with different values. Then, the chosen option must be communicated to downstream regions to drive behavior. The neural circuits involved in value-based decisions are increasingly well understood, but less understood is how decisions shape option representations. We recorded from lateral prefrontal cortex (LPFC), a central hub for transforming options into actions, while non-human primates (NHPs) held two sequentially presented option-value pairs (spatial targets and abstract reward cues) in working memory, then chose the option with the higher value. This revealed how decisions dynamically reorganize option representations in LPFC. We found that, once decisions could be made, representations of chosen and unchosen options rotated into orthogonal subspaces and the chosen option representation was expanded. Before decisions, the first- and second-presented options were maintained separately. After decisions, the chosen option was rotated into a subspace with a consistent representation of the prescribed action, regardless of its presentation order. This suggests a mechanism for value-based decisions where the decision drives a neural subspace reorganization that facilitates selective and efficient readout of the chosen action.

## Introduction

To make an economic choice, we must compare the values of goods available and choose the goods with greater value. The process of making this choice can be broken down into multiple steps including representing available options, assigning values to them, comparing those values, and selecting the option with greater value (1–5). Previous research has studied how values are encoded and compared to make a choice (6–9). But it remains less well understood how multiple options are simultaneously represented in working memory and how these representations are transformed and reorganized by value-based choices.

Lateral prefrontal cortex (LPFC) is a likely candidate site for linking options, values, and actions, and for selecting actions based on their associated values (10–13). Multiple potential options often need to be maintained in working memory during value assignment. The LPFC is also implicated in encoding both choice values and action plans. It receives direct input from the orbitofrontal cortex (14,15), which is involved in processing reward value and other economic factors (1,2,16). It also has dense projections to motor systems (17–19). Neural activity in the LPFC has been shown to reflect the transformation of value into action (11).

Many recent studies have shown that population spiking activity in LPFC (20–29), as well as other frontal areas (30–33), exhibits structured correlations across neurons. As a result, neural activity tends to be restricted to low-dimensional “subspaces” within the high-dimensional space of all possible population activity patterns (reviewed in (34–36)). Structuring activity within such subspaces is thought to flexibly organize information processing (20–25,31,37–40) and communication (41,42).

Motivated by these observations, we recorded from LPFC while non-human primates (NHPs) performed a value-based selection task. To segregate option representations before and after decisions, we used a novel task paradigm that temporally dissociated presentation of options from their value assignment. We examined the representational geometry of the options before and after their relative values were assigned and a decision could be made. We found that, before the decision, the two potential options were held in near-orthogonal subspaces. Once the decision was made, they were transformed to different subspaces that were now orthogonalized by whether their associated actions were chosen vs unchosen, and the representation of the chosen action was expanded. Chosen actions were encoded within the same subspace, regardless of the initial presentation order of their associated option, facilitating their readout to generate behavior. These results show that organized changes in representational geometry of options support value-based decision making.

## Results

### Behavior and analysis

In each trial (Figure 1A), NHPs were presented with a series of two spatial targets (T1 and T2). Each was followed by a reward cue (R1 and R2, respectively) whose color signaled the reward value available if the preceding target was chosen. After the two targets and their values were presented, there was a “Post-decision delay”. The NHPs had all the information needed to decide which target to choose. But they had to wait for a “go” cue (the reappearance of the two targets) before choosing one by saccading to it. Both NHPs most often chose the higher-value target over the lower-value target (97.77% for NHP T; 78.56% for NHP I). Both NHPs reliably chose the higher-value target across all pairs of first and second reward values (Figure 1B; all p<0.001, one-sample t-test). NHP T’s choices were more binary, consistent with their overall higher performance (Figure 1C, top), whereas NHP I’s choices showed a more graded dependence on the relative reward difference (Figure 1C, bottom). Nonetheless, all major effects were robust across both NHPs. Our analyses focus on trials in which NHPs chose the higher-value target.

**Figure 1.**
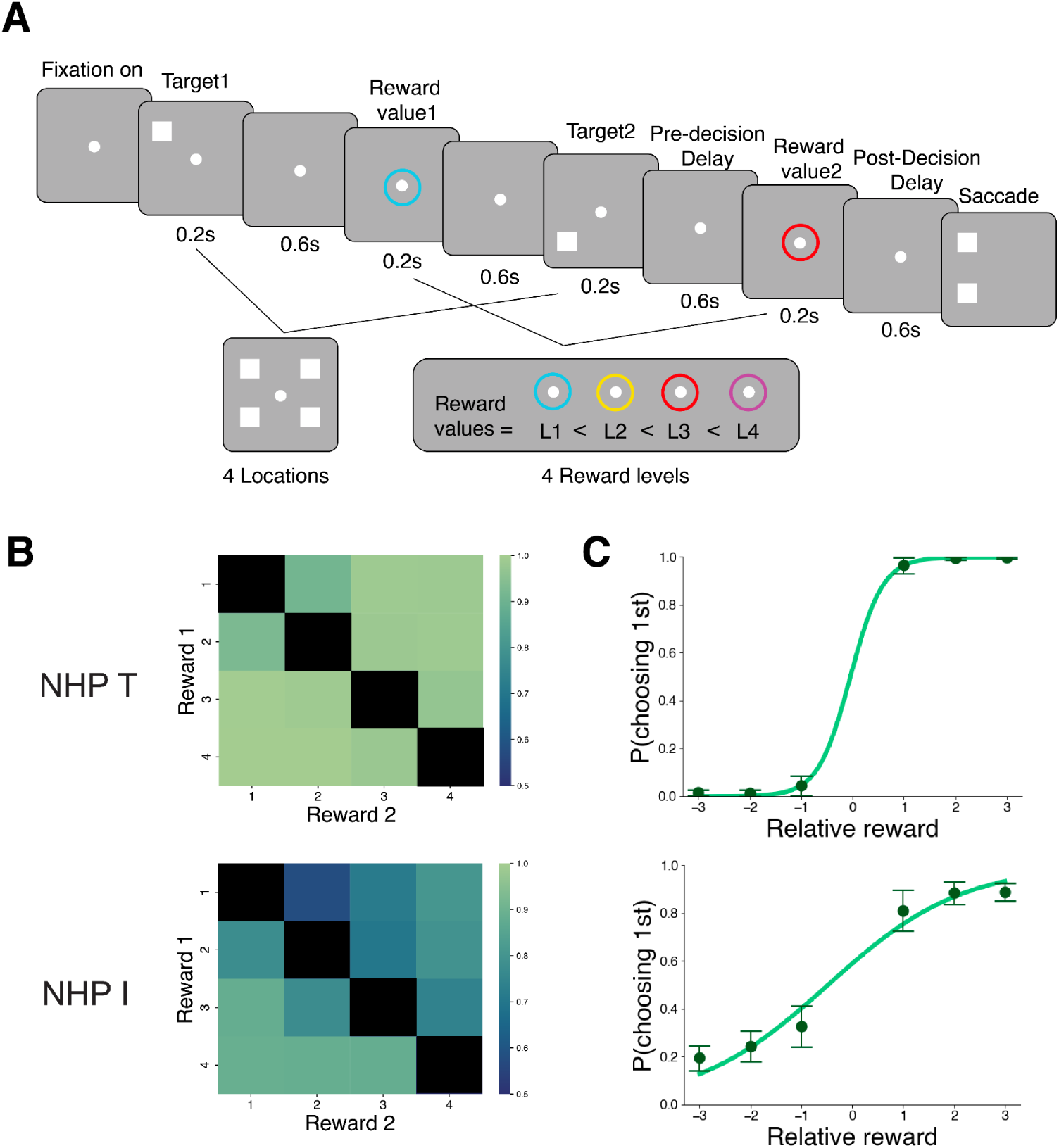
Experimental paradigm and behavioral results. **A** Time course of a task trial, proceeding from left to right. Two spatial targets (T1 and T2) were shown, paired with abstract cues (R1 and R2) signaling the reward value available if the preceding target was chosen. The NHPs’ task was to choose the target associated with the higher reward. **B** Probability of choosing the high-reward target for NHP T (top) and NHP I (bottom) across all combinations of first and second reward values. Values along the diagonal — where the same reward was offered for both targets — were not tested. **C** Probability (mean ± SD across sessions) of choosing the first presented target at different relative reward levels (first-target reward – second-target reward). Both NHPs reliably chose the higher-value target across all pairs of first and second reward values.

Not all trials required an explicit comparison between the reward value of the targets. If the reward value of the first target (R1) was the highest possible value, the NHPs could immediately decide to choose it. Alternatively, if R1 was the lowest possible value, the NHPs could proactively decide to choose the second target when it appeared. We instead focused neurophysiological analysis on the remainder of trials. On those, the NHPs had to wait until the second reward value cue (R2) before they could decide which target had the higher value. Thus, we focused on the time periods just before and after R2. After the second target (T2) but before its value was known (R2), NHPs needed to hold both targets in memory (Pre-decision Delay). Then after R2, they could compare the two targets’ reward values and make a decision (Post-decision Delay). Our data analysis was based on the spiking activity of neurons in LPFC for two NHPs (NHP T: 316 neurons; NHP I: 422 neurons).

### LPFC encoded both reward and target location information

We examined how reward and target information evolved during the decision process. We trained a linear discriminant analysis (LDA) classifier to decode each reward cue’s value and each target’s location from spike rates across the entire neural population. We found an increase of reward information (decoding accuracy) after each reward cue was presented and the reward information was maintained until NHPs made responses (Figure 2A). There was also information about other value-related variables, including chosen value and relative value (Figure S1). Next, we compared location information for chosen vs unchosen targets separately for trials when the first vs second target was chosen. Before the decision, both target locations could be decoded with similar accuracy. But after the second reward cue (R2), when the decision could be made, decoder accuracy for the location of the chosen target increased (Figure 2B,C). Information reflecting the first target location increased when it was chosen (Figure 2B), and there was a corresponding increase for the second target when it was chosen (Figure 2C). This resulted in an interaction between target order and choice (p<0.001, bootstrap test on difference-of-differences; see Methods). We also observed a small subject-specific modulation of this effect that paralleled differences in their behavior (Figure S2). NHP I was slightly biased toward choosing the first target and decoding accuracy difference between chosen and unchosen targets was also slightly greater when the first target was chosen, while choice behavior and neural information were both slightly biased toward the second target for NHP T.

**Figure 2.**
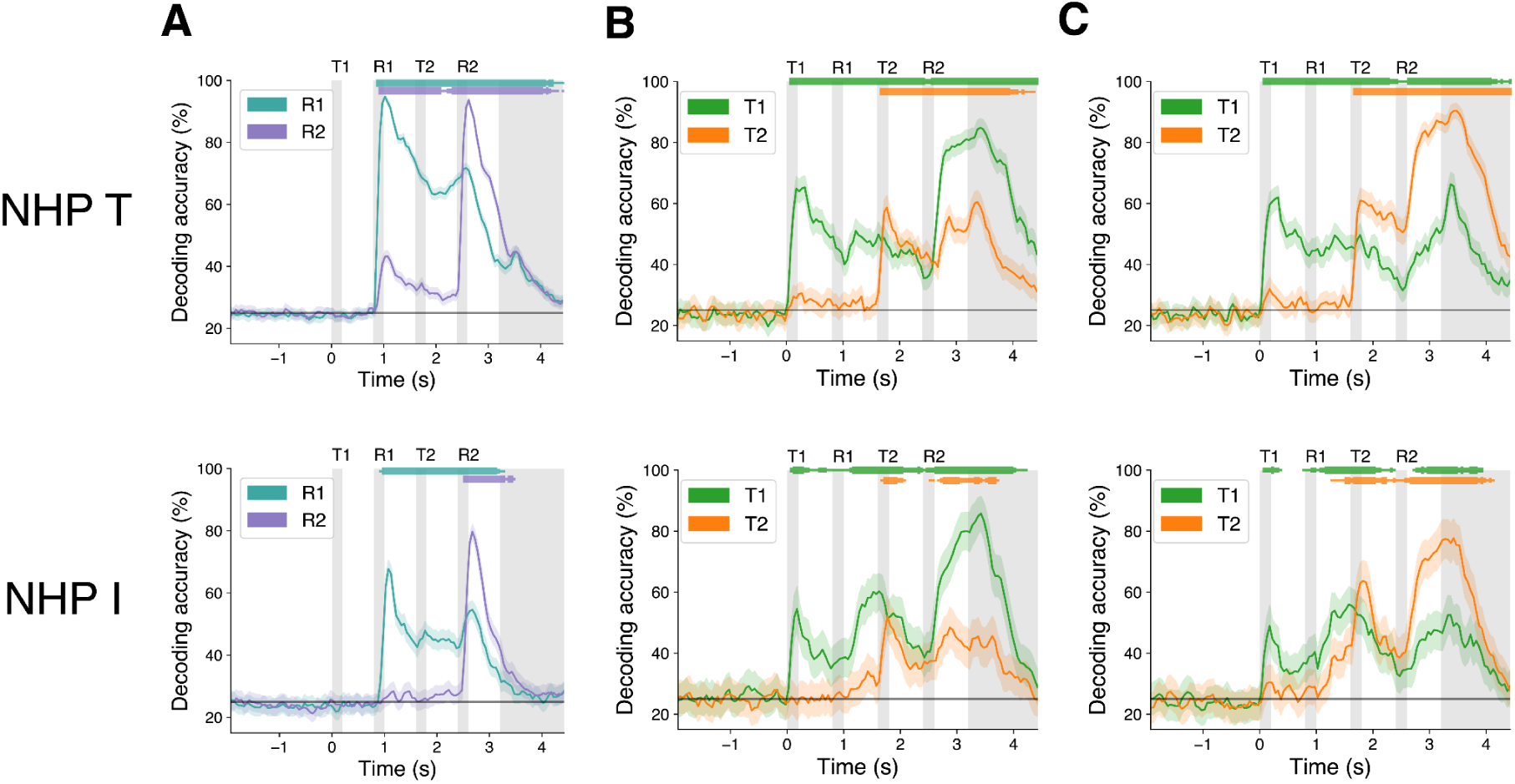
LFP encoded information about both reward cue value and target location. **A** Decoding accuracy of first (R1; cyan) and second (R2; purple) reward cue value. **B, C** Decoding accuracy for the first (T1; green) and second (T2; orange) target location in trials where the first target was chosen (R1>R2; **B**) and where the second target was chosen (R1<R2; **C**) for both NHPs. Width of markers at top indicates significance: p<0.05, p<0.01, p<0.001 for thin, medium, thick markers respectively (corrected one-sided bootstrap against chance level).

The LDA classifier finds the axes in population activity space that maximize the ratio of between-class variance to within-class variance. Thus, the increases in accuracy of target information could reflect either increases in between-class variance or decreases in within-class variance. We found that the between-class variance increased after the decision for both chosen and unchosen targets. This decision-related increase in between-class variance was greater for the chosen targets than the unchosen targets (Figure 3A,C; p<0.001 in both R1>R2 and R1<R2 for both NHPs, chosen vs unchosen targets bootstrap test), similar to the decoding accuracy. By contrast, the within-class variance did not decrease (Figure 3B,D; for NHP T, p=0.403 in R1>R2 and p=0.398 in R1<R2; for NHP I, p=1 in R1>R2 and p=1 in R1<R2; pre- vs post-decision bootstrap test). Thus, the increased decoding accuracy was due to increased separation of the representation of the target locations, not a decrease in their noise.

**Figure 3.**
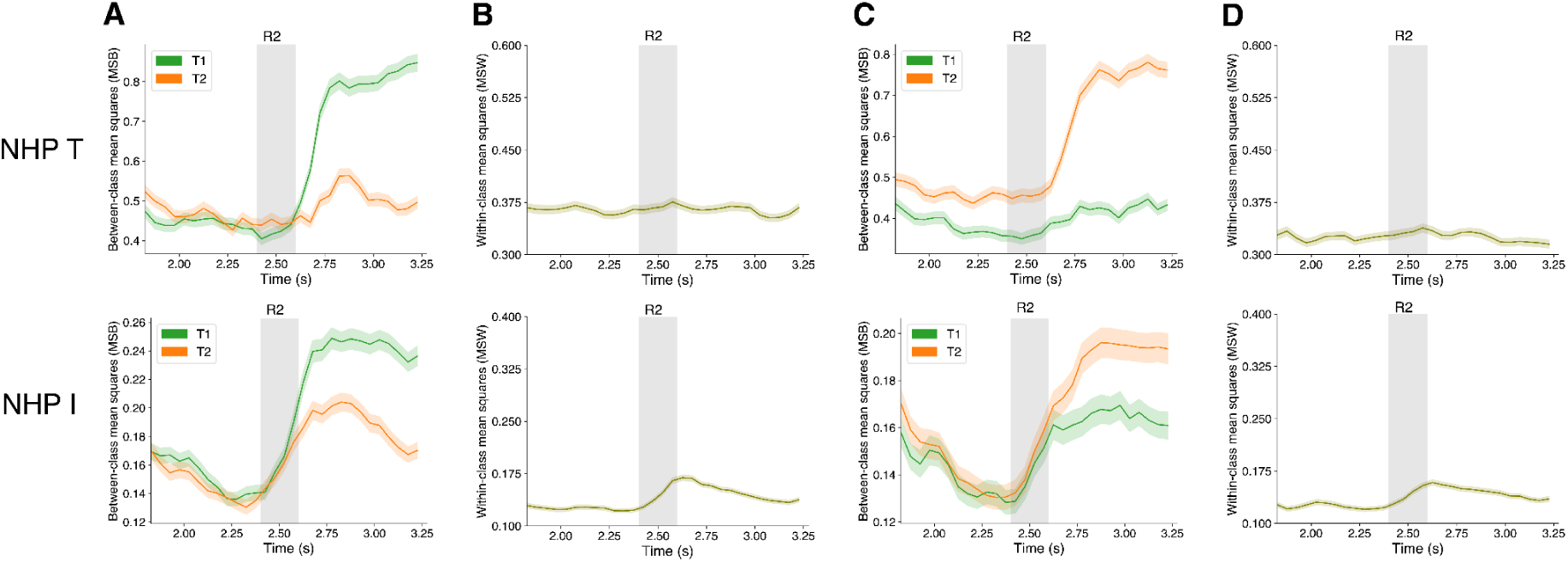
Between- and within-class variance of target information. **A, C** Between-class variance of target locations for the first (green) and second (orange) targets, averaged over all neurons for trials where the first target was chosen (**A**) and where the second target was chosen (**C**). **B, D** Within-class variance of target locations, averaged over all neurons for trials where the first target was chosen (**B**) and where the second target was chosen (**D**). The within-class variance has only a single curve because it was computed for each combination of first and second target locations (i.e. each “cell” in the T1 x T2 design; see Methods). After the decision, the representation of chosen target locations became more separated, resulting in increased decoding accuracy.

### Decisions rotated chosen and unchosen targets into orthogonal subspaces

To examine how decisions transform neural subspaces, we first contrasted the representation of a given target (first/T1 or second/T2) between trials that resulted in different choices. For example, comparing the representation of the first target on trials when it was chosen (R1>R2) vs trials when the *second* target was chosen (R1<R2; an analogous comparison was made for the second target; Figure 4A). To estimate the prefrontal population code for each target, we first fitted a Lasso regression for each neuron’s spike rate using the locations of the two targets as regressors. We then used the fitted regression coefficients across neurons as the population coding vectors for the two target locations (24). Using the fitted regression coefficients, rather than raw firing rates, to estimate population geometry eliminated any biases due to the task design (Figure S3). We visualized the representational geometry of compared target subspaces by projecting the population responses from an example pseudopopulation onto their top three principal components and quantified the alignment between the subspaces. To quantify the alignment between the subspaces in compared conditions, we computed Pearson’s correlation between population coding vectors for corresponding target locations (Figure 4B) (8,43,44). We then averaged correlations across the four target locations to obtain a final “subspace alignment” metric. This measure reflects the geometric alignment of both the compared subspaces and of the arrangement of conditions within each subspace, and it is similar to measures previously used for analogous purposes (8,38). This was performed separately at each time point before and after the decision.

**Figure 4.**
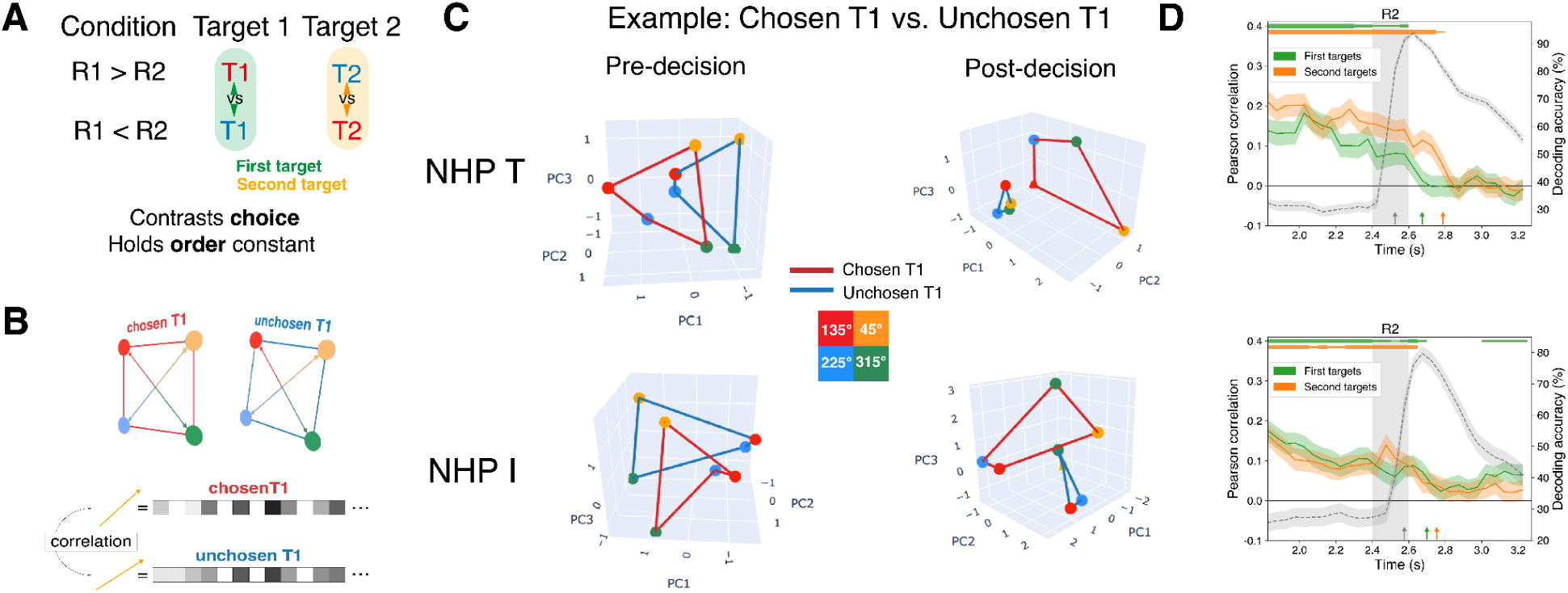
Chosen and unchosen targets became orthogonal after decision. **A** Schematic figure showing the contrast between representations of the same target (first/T1 or second/T2) conditions between trials that resulted in different choices (choose T1 [R1>R2] vs choose T2 [R1<R2]). **B** Subspace alignment between representation of chosen and unchosen first targets (or between chosen and unchosen second targets) was computed as the average Pearson correlation between population coding vectors of corresponding locations between the two targets. **C** Geometry of chosen (red lines) and unchosen (blue lines) first targets before (left) and after (right) decision in reduced 3D space. Four colored dots correspond to the four target locations for each condition. After decision, chosen and unchosen targets are rotated into orthogonal subspaces. **C** Dynamics of subspace alignment (mean ± SD across pseudopopulation bootstraps) between chosen vs unchosen first targets (green line) and between chosen vs unchosen second targets (orange line) across time. The gray dashed line shows information reflecting the second reward value (R2; replotted from Figure 2A), for temporal comparison. Markers at top represent corrected one-sided bootstrap of Pearson correlation for first targets (green) and second targets (orange) against zero. Marker width indicates significance: p<0.05, p<0.01, p<0.001 for thin, medium, thick respectively. Arrows at bottom represent the latency of R2 information (gray arrow) and the decrease in correlation for the first target (green arrows) and for the second target (orange arrow).

We found that decisions resulted in an orthogonalization of the representation of chosen vs unchosen targets. Before decisions could be made (before R2), the subspaces for the first target within trials where it would later be chosen vs unchosen formed parallel planes where the representation of corresponding spatial locations was aligned (Figure 4C, left). After the decision, the subspaces of the chosen and unchosen first targets became near orthogonal (Figure 4C, right). Similar changes in representational geometry were observed when comparing chosen vs unchosen *second* targets (Figure S4).

We quantified this change in population geometry by computing the subspace alignment between chosen and unchosen targets across time before and after the decision (Figure 4D). Alignment was positive before the decision, suggesting that pre-decision representations were similar for targets of the same order. Note that alignment values <1 (perfect correlation) are expected, due to neural noise. Then the alignment significantly decreased (*p*<0.001 for both NHPs, pre- vs post-decision bootstrap test), to a value close to zero. This indicates that chosen and unchosen targets were kept in near-orthogonal subspaces, which minimizes the interference between them. The alignment between subspaces was also quantified using principal angles (45), which showed qualitatively consistent results (Figure S5A). We also computed and compared the latency of value signals and changes in target geometry (Figure 4D, arrows). We found that the value signal preceded the orthogonalization of chosen and unchosen targets, consistent with the geometry change being guided by value-based decisions.

The orthogonal representation of chosen and unchosen targets after the decision could in theory reflect two independent subpopulations of neurons. To test for this, we quantified the overlap between the neural populations with post-decision selectivity for the locations of chosen and unchosen targets (Figure S6A). For NHP T, 32% of neurons selective for the location of the chosen target and 74% of neurons selective for the location of the unchosen target showed selectivity for the locations of *both* targets. For NHP I, 30% of chosen-selective neurons and 74% of unchosen-selective neurons were selective for both targets. Examples of single neurons with distinct spatial tuning profiles for chosen vs. unchosen targets are shown in Figure S6C.This indicates our results reflect different activity patterns for chosen and unchosen targets across a largely overlapping neural population, rather than separate subpopulations.

### Decisions brought orthogonal subspaces into alignment for chosen targets

Next, we contrasted representations of conditions where targets had different temporal order but resulted in the same choice (Figure 5A). That is, we computed the alignment between the subspaces of the first and second targets on trials where each of them were *chosen*, and analogous comparisons between the first and second targets on trials where they were each *unchosen* (Figure 5B). Before the decision, the first (solid lines) and second (dashed lines) targets each formed approximately planar structures that were nearly orthogonal (Figure 5C, left). After the decision, the chosen first and second targets (derived from separate trials) formed parallel, aligned planes (Figure 5C, right). The alignment between the subspaces of the two targets before the decision was slightly positive but close to zero (Figure 5D). This indicates that the population activity patterns coding for the same set of target locations were initially nearly orthogonal for the first vs. second targets. After the decision, the alignment between the chosen first and second target subspaces (derived from separate trials; Figure 5D, red lines) increased significantly (*p*<0.001 for both NHPs, pre- vs post-decision bootstrap test). In contrast, the alignment between the first and second targets when they were *unchosen* (Figure 5D, blue lines) increased only slightly for one NHP (NHP T; p<0.001) and showed no change for the other NHP (NHP I; p=0.866). The alignment between subspaces was also quantified using principal angles, which showed qualitatively consistent results (Figure S5B). Again, the alignment between chosen targets followed the onset of the reward value signal, consistent with reward-based decision driving the alignment (Figure 5D, arrows).

**Figure 5.**
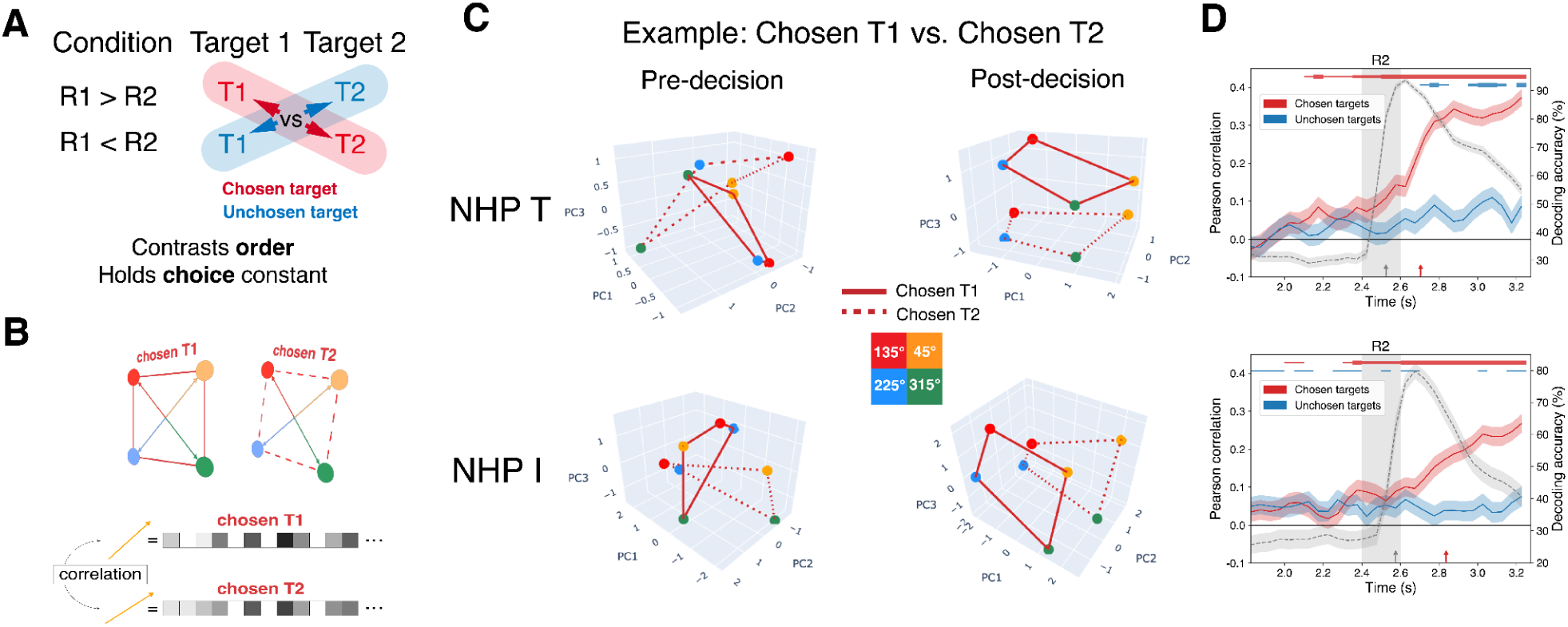
Target representations were reorganized by choice after decisions. **A** Schematic figure showing the contrast between targets with different order of presentation but resulting in the same choice. **B** Subspace alignment between first and second targets when they were each chosen (or between the two targets when they were both unchosen) was computed as the average Pearson correlation between population coding vectors of corresponding locations across the two targets. **C** Geometry of the first (solid lines) and second (dashed lines) targets when they were chosen, before (left) and after (right) decision in reduced 3D space. Four colored dots correspond to the four target locations. The first and second targets are initially held in orthogonal subspaces. After the decision, the first and second target are rotated into the same, aligned subspace, but only when they are chosen (on separate trials). **D** Dynamics of subspace alignment (mean ± SD across pseudopopulation bootstraps) between first and second targets when they were each chosen (red line) and when they were each unchosen (blue line) across time for both NHPs. The gray dashed line represents the R2 information (replotted from Figure 2A). The markers at top represent corrected one-sided bootstrap of Pearson correlation for chosen targets (red) and unchosen targets (blue) against zero. Marker width indicates significance: p<0.05, p<0.01, p<0.001 for thin, medium, thick respectively. Arrows at bottom represent the latency of R2 information (grey arrow) and of the alignment between chosen targets (red arrows).

The observed alignment of chosen target representations could allow downstream areas to read out the location of the chosen target with a single decoder, regardless of its initial presentation order. We tested this by training classifiers to decode the chosen target location on trials where the *first* target was chosen, then testing on trials where the *second* target was chosen, and vice versa (Figure S7). This cross-decoding was nearly as accurate as training and testing on the same target-choice condition, confirming the feasibility of readout using a single decoder. For the orthogonal subspaces before decisions, we confirmed that they are supported by an overlapping neural population selective for both first and second targets, rather than separate subpopulations (Figure S6B). For NHP T, 60% of neurons selective for T1 and 55% of neurons selective for T2 showed mixed selectivity for both targets. For NHP I, 59% of T1 neurons and 60% of T2 neurons showed mixed selectivity for both targets. Examples of single neurons with distinct spatial tuning profiles for first vs. second targets are shown in Figure S6D.

Before the decision, the LFPC population representation was organized in a format that kept the representations of the first and second targets independent. After the decision, it reorganized into a format in which the chosen location could be read out consistently, independent of whether it was presented first or second. Thus, the two contrasts (Figure 4 and Figure 5) combined indicated a reorganization of target representation from order of presentation to choice.

### Aligned target subspaces remained separable by choice

The alignment between chosen target subspaces could in theory reflect a transition to a purely spatial motor preparation code. Instead we found that post-decision population activity also maintained an orthogonalized representation of which target was chosen. We trained an LDA classifier to decode choice information (whether the first or second target was chosen on each trial). We found that choice information increased after the second reward cue, and remained strong through the post-decision delay (Figure 6A). We confirmed this by projecting the population coding vectors of chosen targets onto the LDA-estimated choice axis. We found that the two chosen targets could be separated on the choice axis after R2 and remained separable throughout the post-decision delay (Figure 6B). The choice axis was orthogonal to both aligned target location subspaces (Figure 6C). These results are also consistent with the example post-decision population geometries shown in Figure 5C. These results suggest a post-decision transformation of population geometry that maintained target selection information in a format independent of the selected location, rather than a transition to a pure spatial motor-preparation signal.

**Figure 6.**
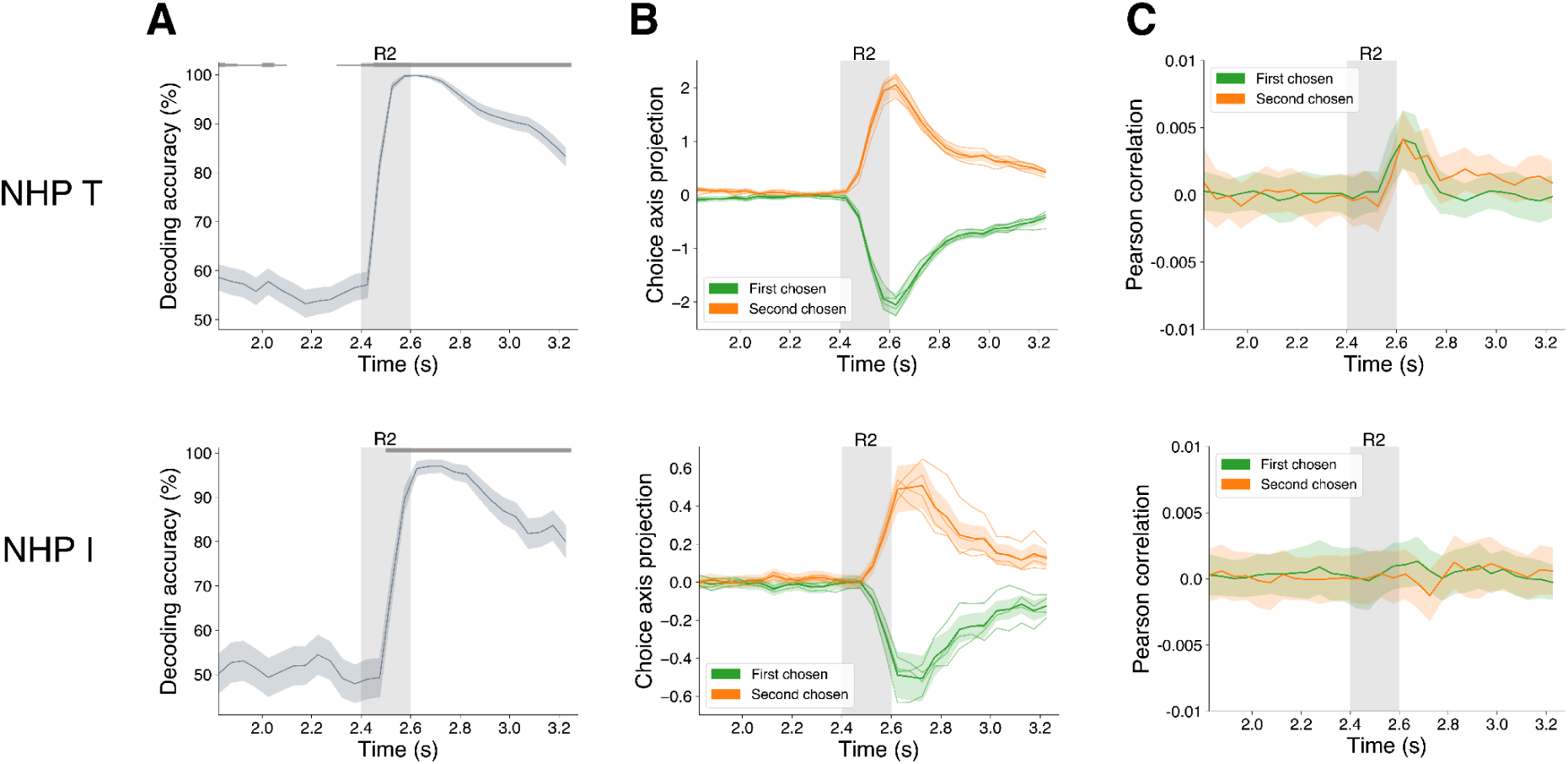
Aligned target subspaces remained separable by choice. **A** Decoding accuracy of choice (whether the first or second target was chosen) before and after decision for both NHPs. The markers at top represent corrected one-sided bootstrap of decoding accuracy of choice against chance level. **B** The mean projection of population coding vectors of chosen targets (first target chosen: green line; second target chosen: orange line) onto the decoder choice axis. Lighter lines show the projection of each individual location for each chosen target. **C** Subspace alignment between choice axis and target location axes. Green line represents the alignment between choice axis and the first target chosen and orange line represents the alignment between choice axis and the second target chosen.

### Early subspace reorganization reflected early decisions

For the findings presented so far, we excluded the trials where the first reward cue (R1) signaled the highest or lowest possible value. In these trials, the NHPs could assume the second target’s reward must be lower or higher in value, respectively, than the first target’s reward. Thus, they could make an early decision without waiting for the second reward cue (R2). Here, we examined these previously excluded “early decision” trials. We hypothesized that if an NHP was able to reliably make an early decision, the observed decision-related changes of neural representation (increased information of chosen target and subspace reorganization) would also appear earlier in the trial than in the previously considered “late decision” trials.

We found mixed evidence for early decisions in the neural representations. For NHP T, information about the first target and second target increased when the first reward cue signaled the highest and lowest possible value respectively (Figure S8A, solid green line; Figure S8B, solid orange line). NHP I showed a similar, but non-significant, trend only in trials where the first reward cue signaled the highest value (Figure S8C). The weaker and asymmetrical results in NHP I may be related to differences in their behavior between these conditions. When the first target had the highest possible value, NHP I performed well, regardless of the second value. When the first target had the lowest possible value, performance was poorer, especially when the second target was similar in value (Figure 1B). Thus, behavioral and neural results both suggest this subject may have made early decisions only when the first value was highest. These results offer some evidence that, when the NHPs can make early decisions, the increased information of chosen targets also occurs correspondingly early.

We also found evidence for early decisions in the decision-related neural subspace reorganization (orthogonalization and alignment; Figure S9). For NHP T, the early decision trials resulted in earlier orthogonalization of chosen vs unchosen targets earlier relative to late-decision trials (Figure S9B,C, left), similar to the early changes in decoding accuracy. Similar, but weaker effects were observed in NHP I (Figure S9B,C, right). For both NHPs, there was also some evidence for early alignment of chosen target subspaces regardless of the initial target presentation order on early-decision trials (Figure S9E). Also, the exact value of first reward (R1) had a graded effect on the alignment between first targets, such that targets with more similar values were more aligned (Figure S10). Taken together, these results provide evidence that, when early decisions are possible, the neural subspace reorganization reflecting decisions can happen correspondingly early.

## Discussion

We found that, during value-based decision making, LPFC encoded both reward and option (i.e. target location) information and the representational geometry of options changed dynamically as the decision unfolded. When two potential options were simultaneously held in working memory, they were held in separate, near-orthogonal subspaces corresponding to their order of presentation. This presumably allowed the brain to keep the options separated by their behavioral context (i.e., order of presentation) before their relative values were known. After the choice could be made, neural subspaces of the prescribed actions rotated such that the representation of the chosen actions was aligned, regardless of their order of presentation. This shared subspace of chosen actions was near orthogonal to the subspaces of both unchosen actions, reducing interference from irrelevant information. In addition, options associated with higher values showed increased feature separation, which likely produces more accurate readout of the chosen action in downstream motor areas.

Orthogonalization of neural representations has been suggested as a mechanism to keep information independent (22,23,25,31,37–40,46–49). Neural activity changes reflecting one type of information would have no effect on other information held in an orthogonal subspace. Downstream processes could read out one with no interference from the other. Evidence has suggested mutual orthogonalization of sensory inputs and working memory (37), of working memory sequences (23), of perceptual choice and motor plans (50), and of reach preparation and movement (31). Our results provide converging evidence for this idea in the context of two competing decision options. Orthogonalizing the two targets by order of presentation before the decision could keep their representations independent until it was time to select one. It may also support binding of the targets to their subsequent reward value assignments (51). After the decision, keeping the representations of chosen and unchosen targets orthogonal may allow readout of the chosen target location without interference from the unchosen target.

Subspace alignment has been suggested to be important for discarding irrelevant information, while bringing relevant information into a common format to facilitate downstream readout (44,52–56). This could allow readout using a single linear decoder, regardless of any information irrelevant to the readout. In our study, the post-decision alignment of chosen targets onto a common subspace allows readout of target location using a single decoder regardless of their initial temporal order. This might, in part, reflect a transition to a neural code for motor preparation. However, importantly, information about the temporal order of the chosen target did not simply disappear. Instead, it was robustly maintained through the entire trial, but along a population axis orthogonal to the aligned subspace coding for chosen target location. This suggests our results are better described by a transformation of population geometry that rotated the selected target into a common format suitable for downstream readout, rather than simply reflecting choice-independent preparatory motor activity. Preserving choice information in an orthogonal axis may enable LPFC to track the decision history and support later credit assignment without interfering with the readout of motor output (57).

The geometric changes we report largely reflect effects at the population level that are not easily reducible to effects at the single-neuron level. We found a reorganization from orthogonalization by target temporal order (first vs second) to orthogonalization by choice status (chosen vs unchosen). In both cases, the orthogonal subspaces largely reflected different patterns of activation across overlapping neural populations, rather than activation of separate subpopulations. This converges with analogous results across many different domains (38,58,59). It is also consistent with the well-established “mixed selectivity” of PFC neurons, where multiple task variables are encoded within overlapping neural populations and reorganized through changes in population activity patterns (26,44,60). In contrast, the increased population separation of chosen target locations could be explained by a gain increase at the single-neuron level. This is consistent with a large body of work showing enhanced single-neuron spiking activity and information content for behaviorally relevant information (61–67). However, we (and others (35,68,69)) argue that, even in this case, population geometry provides a concise unifying analytical framework for understanding diverse neural effects, which likely better approximates the level at which neural computation and communication actually happens in the brain.

Making decisions requires integrating various feature dimensions of each option into a representation of overall value, choosing the option with the highest value, then mapping the chosen option onto an appropriate action (1,2,70). Different brain regions have been implicated in different stages of this decision process. Evidence suggests orbital (OFC) and ventromedial PFC (vmPFC) encode an abstracted representation of overall option value and compare values of different options (9,71–77). Anterior cingulate cortex and motor-related areas such as supplementary eye field (SEF) engage in action selection and encode the chosen action in economic choices (78–80). A key region that can use the value signals to perform option selection and inform the action mapping is LPFC, which is involved in option selection and the mapping from chosen options onto appropriate actions (11,81,82).

Our results showed that LPFC may be involved in multiple stages of this decision process. First, LPFC holds multiple options online before decisions, within orthogonal subspaces. This resembles findings in orbital and medial PFC, where option values are encoded in orthogonal subspaces (51). Second, LPFC induces a rotation in LPFC subspaces representing potential options and/or their associated actions, guided by the emerging value-based decisions. This is analogous to the choice-induced rotation of the OFC/vmPFC subspaces representing option values induced (58). We further show the subspace rotation reflects a shift from organization by offer presentation order (first vs second) to organization by choice status (chosen vs unchosen). Previous work has shown that value signals can guide action selection by modulating the buildup of choice-related activity in ACC (80). Our results provide a complementary perspective by showing that value-driven selection also reshapes the geometry of option subspaces, reorganizing subspaces from an order-based format into a choice-based format that supports action selection. Moreover, LPFC does not induce the decision-related changes in neural representation only after both values were revealed. Instead, it shapes target representations using the partial value information even before both values were revealed. Target information was enhanced when it was associated with highest value, and targets of more similar values occupied more aligned subspaces. This suggested that partial value information can begin to bias target representations before the full decision is available, thereby facilitating subsequent selection.

In summary, our results illustrate the dynamic subspace reorganization supporting option maintenance and selection in economic decisions. Before making the decisions, the brain maintained two possible options using orthogonalization. When it came time to make a decision, the decision induced a rotation of the chosen options into a common format, facilitating their translation into action. These findings highlight a role for dynamic changes in representational geometry in working memory and decision making.

## Methods

### Non-human primate (NHP) subjects

Two adult macaque monkeys (*Macaca mulatta*), monkey T (male, 10 years old, 13 kg) and monkey I (female, 15 years old, 9 kg) served as subjects in this study. All procedures followed the guidelines of the Massachusetts Institute of Technology Committee on Animal Care and the National Institutes of Health.

### Task paradigm

We trained the NHP subjects on a value-based selection task (Fig. 1A). In each trial, a sequence of two spatial targets (white squares; 200 ms each) appeared at distinct locations (four possible at polar angles of 45°, 135°, 225°, and 315°, all at 5° eccentricity). All 12 possible distinct first/second target location combinations were presented with equal probability. Following each spatial target (after a 600 ms delay), an abstract reward cue (200 ms) signaled the magnitude of juice reward that would be obtained by remembering and later directing a response toward the immediately preceding spatial target. The reward cues were non-spatially-informative annuli, with four possible colors reflecting distinct reward values offered. The colors of the four reward cues differed between subjects, to disentangle low-level visual features from the associated reward offer. The associated absolute reward values were also slightly different between subjects, to maintain comparable behavior. For NHP T, blue=1 juice drop, yellow=5 drops, red=8 drops, and magenta=10 drops. For NHP I, yellow=1 juice drop, green=6 drops, blue=9 drops, and cyan=13 drops. Each trial always contained two distinct spatial targets and two distinct reward cues—there were never ambiguous trials where the first and second cue signaled the same reward or where the first and second spatial locations were identical. Otherwise, spatial locations and reward cue values were set randomly and independently on each trial—any location could be paired with any offered reward value. Finally, response targets were displayed at the locations of both spatial targets from that trial. Subjects were allowed to choose one by making a saccadic response to either location and received the reward signaled by its associated reward cue. Note that either choice of response target was rewarded. So “correct choice” in this task means that the subject chose the target with the larger reward.

### Electrophysiological recording

Recordings were conducted with multiple single-contact tungsten electrodes (FHC). Raw analog data was amplified and recorded at 30 kHz using the Blackrock Cerebus system. Spikes were extracted by band-pass filtering (250–7500 Hz) and manually setting online amplitude thresholds. Spikes were sorted manually offline into isolated single units (Plexon OfflineSorter). All units were included in analyses and no other preprocessing was performed on the spiking data. Behavior was controlled using the PsychToolbox (monkey T) or MonkeyLogic (monkey I) systems, and eye position was monitored at 1 kHz (Eyelink 1000).

Recordings were unilateral and spanned both ventrolateral and dorsolateral PFC (Brodman areas 46, 45, and 8A; Figure S11). Frontal Eye Field (FEF) sites were localized using standard microstimulation mapping—sites near the arcuate sulcus where low-current (<50 μA) stimulation reliably evoked saccades. Removing the few putative FEF and premotor cortex sites produced nearly identical results (Figure S12).

The data includes 23 sessions from NHP T and 17 sessions for NHP I. Each session, we recorded from an average of 24±2 (mean±SD) electrodes for NHP T and 33±10 electrodes for NHP I. This resulted in a total of 316 neurons from NHP T and 422 neurons from NHP I.

### Behavioral analysis

All analyses described here and below were performed with custom codes written in Python. We computed the behavioral accuracy for each of the 12 combinations of the first and second reward offer. Note that there are only 12 reward combinations because the same reward cue was not repeated twice within a single trial.

To estimate the influence of reward difference on accuracy, we computed the probability of choosing the first spatial target as a function of reward difference (reward 1 – reward 2; resulting in six levels of reward difference = {–3,–2,–1,+1,+2,+3}) and we fitted the accuracy with a logistic function. Behavioral results (Figure 1B,C) are presented as means ± standard deviations across all sessions for each subject. Their significance was quantified using sessions as observations (*n* = 23 sessions for NHP T, 17 sessions for NHP I).

### Neural data processing

All neural data analyses used only correct trials (where the subject chose the higher value target) and only sessions with at least 15 correct trials for each condition (target location 1 ⨉ target location 2 ⨉ choice) were included. For most analyses, we also only included trials in which the first reward does not have the highest or lowest values. In trials where the first reward had the highest or lowest value, subjects could immediately infer which target to choose, without making any comparison to the second reward value. Thus, we included the trials in which the first reward has intermediate reward levels and focused the analysis on the delay period before and after the second reward was presented, when a decision could be made.

For most neural analyses, we generated pseudo-simultaneous population responses (“pseudopopulations”) by combining neurons across sessions as if they were sampled simultaneously, subsampling trials to match their count for each condition across neurons (83). We randomly resampled pseudopopulation trials 1000 times to estimate means and standard deviations of summary statistics, and to evaluate their statistical significance. We calculated spiking rates using a 150 ms sliding window with 50 ms step size.

### State-space analysis

#### Linear regression

To deal with the non-independence between the two target locations in our task design (i.e. the two spatial targets cannot be the same within a single trial), we used linear multiple regression to quantify how each neuron’s activity was independently affected by the two spatial targets, and used the resulting fitted coefficients as our estimate of the neural population response (24). We assumed that the average spiking rate *y_i_*_,*t*_of the *i_t_*_ℎ_ neuron at time *t* is the linear combination of effects due to the first and second spatial targets, such that:

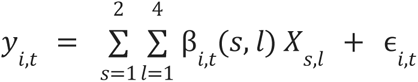

Where β(*s*, *l*) is the coefficient for location *l* (within the set {45°,135°,225°,315°}) of target *s* (within the set {T1,T2}), *X_s_*_,*l*_ is the regressor reflecting location *l* of target *s* (“dummy” encoded as {0,1}), and ɛ*_i_* is the residual error. For each trial, a sequence of two targets can be represented as an 8-dimensional vector and each target location is one-hot encoded. For example, sequence [45°, 135°] can be represented as [(1 0 0 0), (0 1 0 0)]^T^. We wanted to estimate coefficients for all possible location conditions, but this naturally resulted in an over-determined, rank-deficient design matrix. To solve rank deficiency and avoid overfitting, a Lasso regularization term was added to the linear regression model, which has previously been successfully used for analogous analytical situations (24). The regularization amplitude *α* was selected using 5-fold cross-validation. We fitted Lasso models across a grid of candidate *α* values ranging from 1 to 10⁻⁵ and selected the *α* that minimized the cross-validated prediction error. This Lasso regression method successfully corrected for any bias due to the non-independent design, both for simulated data with known ground truth and for real data from example neurons (Figure S3), as expected from previous work (24).

At each time *t*, we concatenated the coefficients for location *l* of target *s* across neurons and we defined this vector of coefficients as the population response of location *l* of target *s* at time *t*. We separated trials based on whether the first reward was higher (first target chosen) or whether the second reward was higher (second target chosen) and performed separate linear regressions for these two conditions. Using the first-target-chosen trials for regression, the population response of the first target represents a chosen target, while the population response of the second target represents an unchosen target. Using the second-target-chosen trials for regression, the population response of the first target represents an unchosen target, while the population response of the second target represents a chosen target.

#### Subspace alignment

We computed the degree of alignment between the representations of two targets by computing the Pearson’s correlation between the mean population response vectors for the corresponding locations of the two targets (e.g. location 1 of the first target and location 1 of the second target) and averaging across the four locations. We refer to the resulting index as the “subspace alignment” between two representations.

Alignment between subspaces was also quantified using principal angles (45). We estimated a 2D subspace for each target representation by applying principal component analysis (PCA) to the condition-mean population activity and finding the first two principal components. We computed the principal angles between the two subspaces by performing singular value decomposition on the product of the two subspaces. The singular values correspond to the cosines of the principal angles between the subspaces. We report the smallest principal angle as a summary measure of subspace alignment, with smaller angles indicating stronger alignment between the two target representations.

Two types of contrasts were made between targets. First, we compared conditions with the same target presentation order, but resulting in different choices (Figure 2). That is, we computed the subspace alignment between the first target when it was chosen vs unchosen, and between the second target when it was chosen vs unchosen. Second, we compared conditions with different target order, but resulting in the same choice. That is, we computed the subspace alignment between the first target when it was chosen vs the second target when it was chosen, and between the first target when it was unchosen and the second target when it was unchosen.

### Decoding analysis

We trained linear discriminant analysis (LDA) classifiers to decode the reward values, value-related variables (chosen value and relative value), and the location of the first and second spatial targets, separately within each time window. To equate the number of trials for each class, trials were randomly subsampled to match the class with the lowest trial count. For decoding the chosen reward value (Figure S1A), we excluded the lowest reward value, since in correct trials it was never chosen, and decoded the three remaining reward values. Within each of the 1000 randomly resampled pseudopopulation, we performed 5-fold cross-validation to compute the decoding accuracy. Before training the decoder at each fold, we z-scored the training and testing data for each neuron based on the mean and standard deviation across all trials of the training data. To reveal how decisions affect the target representations, we trained the classifier separately for trials in which the first reward was higher and for trials in which the second reward was higher. To examine whether the observed decision-related improvement in decoding accuracy was caused by increased between-class variance or decreased within-class variance, we computed between-class and within-class variance. Between-class variance for a spatial target is defined as the sum of squared differences between each location mean and the overall mean divided by (number of locations - 1). We quantified within-class variance as the variation around the mean neural response for each combination of the first and second target locations. In this case, within-class variance is defined as the sum of squared difference between individual response and the mean response in each location combination divided by (total number of neural responses - the total number of location combinations).

### Single-neuron selectivity and population overlap analysis

To quantify single-neuron selectivity for each target, we pooled each neuron’s spiking activity within the pre-decision or post-decision epoch and fitted a linear regression model in which the two target identities were included as categorical predictors, each with four possible levels. For each neuron and each target variable, the first level was omitted as the reference category, and an intercept term was included in the model. Thus, regression coefficients for the remaining levels quantified changes in firing rate relative to the reference level. A neuron was classified as selective for a given target if at least one regression coefficient associated with that target variable was statistically significant (*p* < 0.05; two-tailed t test).

### Quantification and statistical analysis

Unless otherwise noted, bootstrap-like resampling of pseudopopulation trials was used for all error quantification and statistical testing (84). Plots show the mean and standard deviation of a measure (e.g. subspace alignment, decoding accuracy) across 1000 random bootstrap-like resamples of trials to form a pseudopopulation, as detailed above. The standard deviation of a summary statistic across bootstraps is an estimate of the standard error of the statistic (84). Thus, plotted error bars represent a bootstrap estimate of the standard error of the mean. For statistical testing, *p* values were generated by counting the proportion of bootstrap samples less than the value of a statistic expected under the null distribution (one-sided test that statistic is greater than expected by chance). For comparisons before vs after decisions, the statistic was the paired difference between values before (1.8–2.4s) and after (2.6–3.2s) the decision point (R2). To evaluate the target order ✕ choice interaction effect on decoding accuracy (Figure 4A,B), we computed a “difference-of-differences” statistic commonly used for bootstrap tests of interactions (85,86). For each bootstrap sample, we computed the difference in decoding accuracy between the two target orders (T1 – T2) and between the two choice conditions (choose T1 – choose T2), and then computed the difference of these differences ((T1 – T2) – (choose T1 – choose T2)). The distribution of this difference-of-differences statistic across bootstrap resamples was evaluated against a null value of 0. For statistics evaluated at multiple individual timepoints, *p* values were corrected for multiple comparisons across timepoints using the Benjamini/Hochberg procedure for controlling the false-discovery rate (87).

## Acknowledgements

We thank Will Dorrell, Mikael Lundqvist, and Jonatan Nordmark for helpful comments and discussions on this work. This work was supported by Office of Naval Research MURI N00014-23-1-2768 (E.K.M.), Army Research Office W911NF2410228 (E.K.M.), NEI 1R01EY033430-01A1 (E.K.M.), The Freedom Together Foundation (E.K.M.), The Picower Institute for Learning and Memory (E.K.M.), NINDS Sustainable Transformation of Institutional Research Rigor (STIRR) Program (H.L.), and the Knut and Alice Wallenberg Foundation (N.C.).

## Author contributions

J.R., and E.K.M. designed the experiments. J.R. collected the data. S.L.B. curated the data. H.L., N.C., and S.L.B. designed and carried out the data analysis. H.L., N.C., S.L.B., and E.K.M. wrote the manuscript, and E.K.M. supervised the study.

## Resource availability

Requests for further information and resources should be directed to and will be fulfilled by the lead contact, Earl K. Miller

## Supplementary figures

**Supplementary Figure S1.**
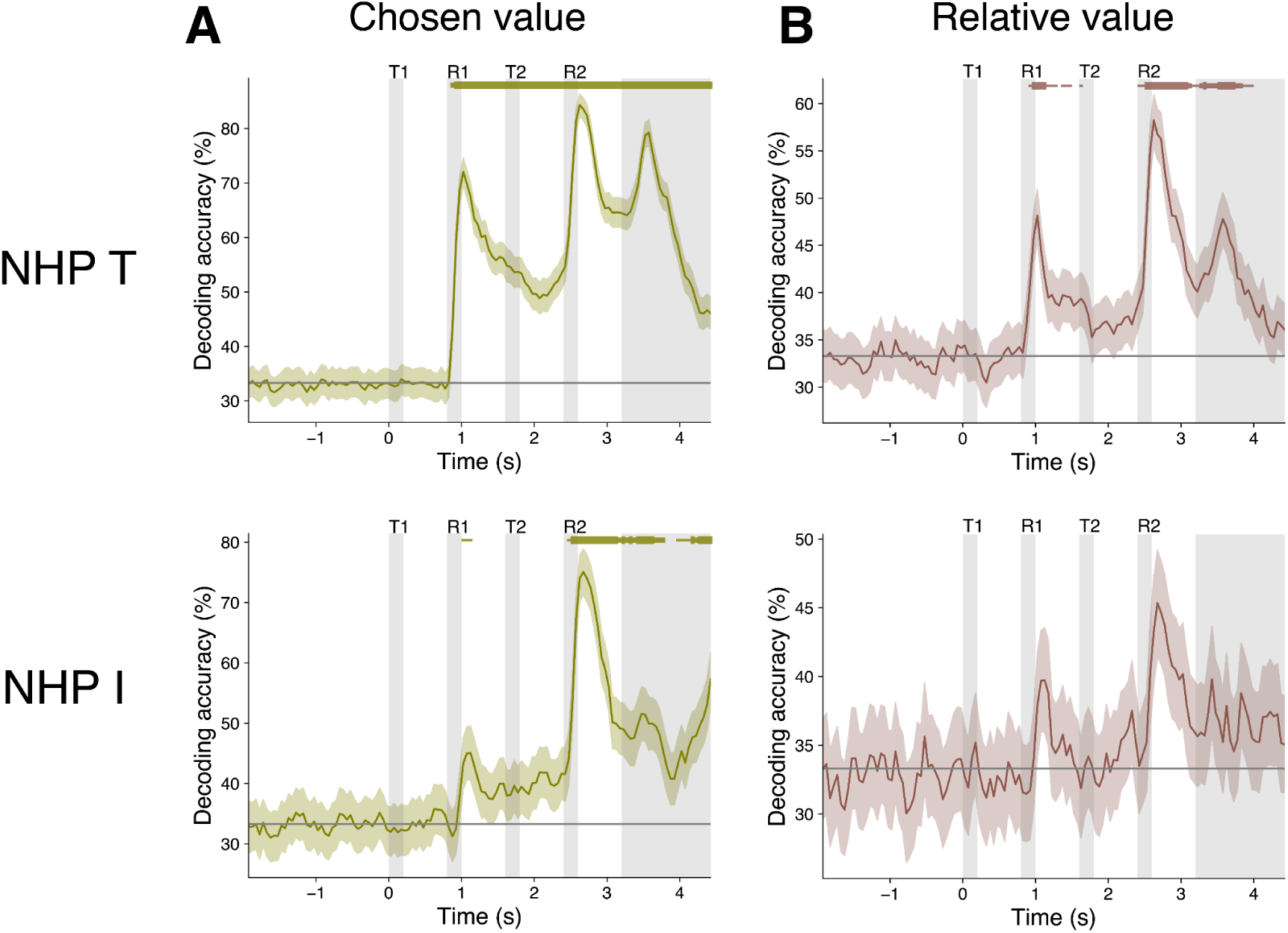
LPFC conveyed information about multiple value-related variables. **A, B** Decoding accuracy of chosen value (**A**) and relative value difference (R2–R1; **B**) for both NHPs. Width of markers at top indicates significance: p<0.05, p<0.01, p<0.001 for thin, medium, thick markers respectively (corrected one-sided bootstrap against chance level).

**Supplementary Figure S2.**
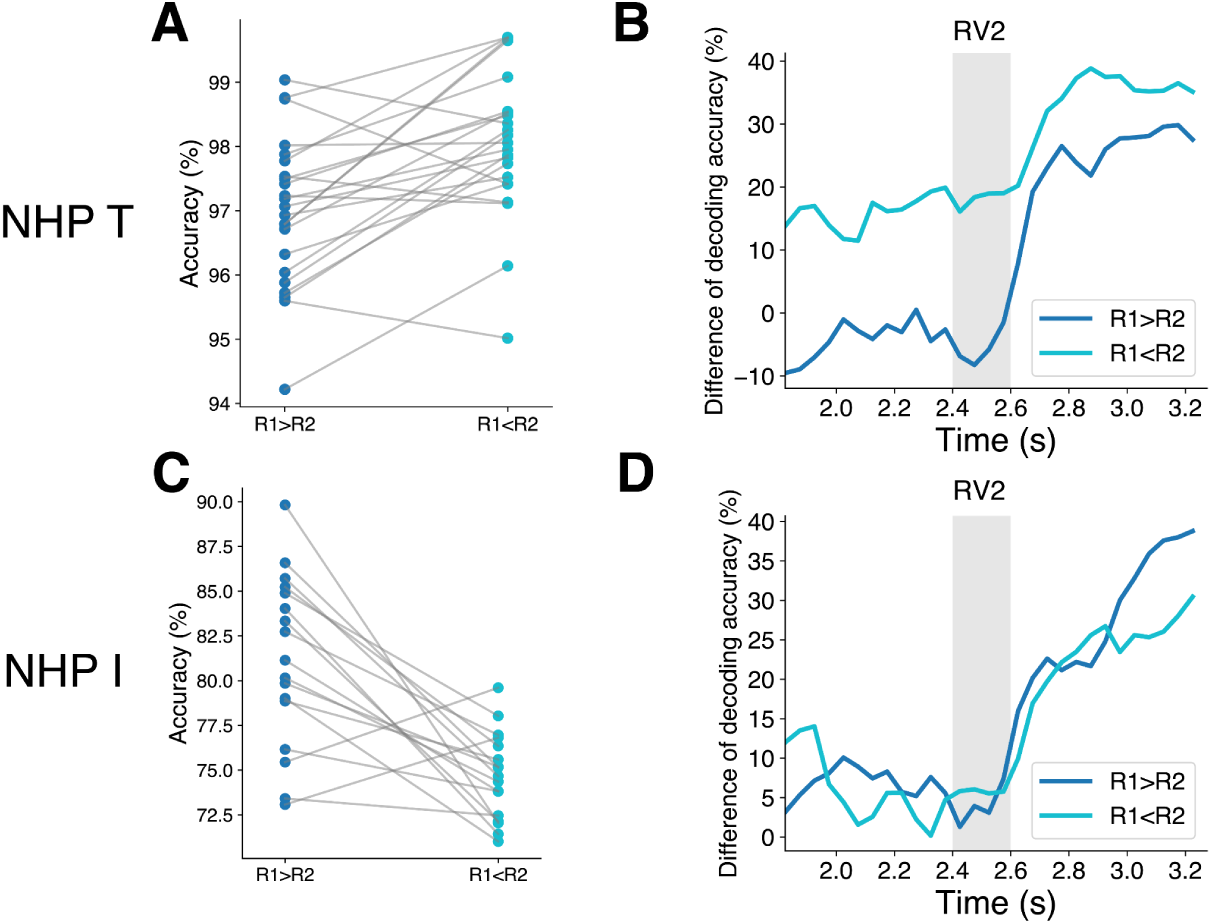
Behavioral bias and the neural representations that reflect the behavioral bias. **A,C** Behavioral accuracy within each session (dots) of correctly choosing the higher-value target when the first target had higher value (R1>R2; blue) and when the second target had higher value (R1<R2; cyan). **B,D** Difference of neural decoding accuracy between chosen and unchosen target for condition R1>R2 (blue) and R1<R2 (cyan).

**Supplementary Figure S3.**
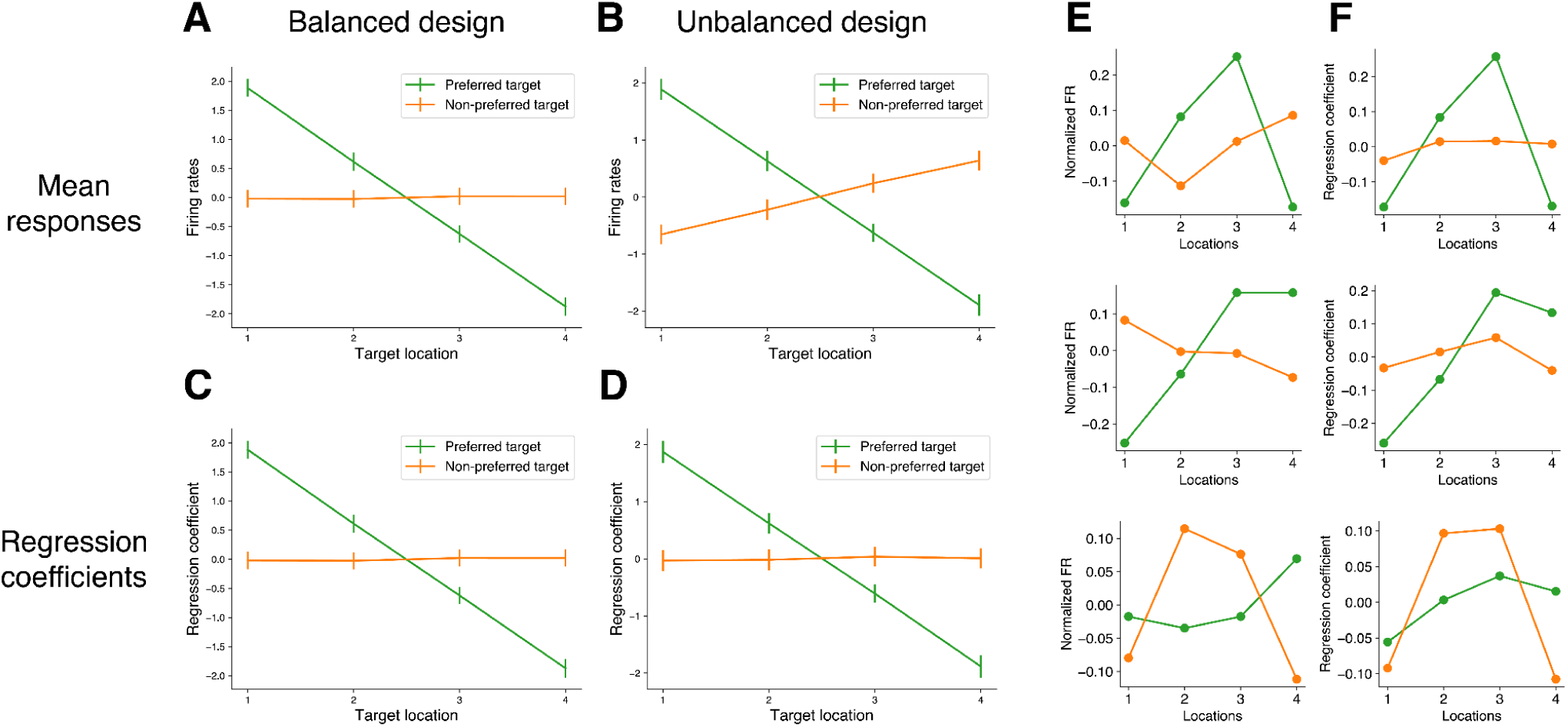
Lasso regression removed anti-correlation bias introduced by non-independent experimental design. **A,B** Mean firing rates across target locations for a simulated neuron with known ground-truth spatial tuning for target 1 (green), but not for target 2 (orange). Under a fully balanced task design (**A**), no tuning is (correctly) observed for target 2. A non-independent design (**B**), where the same location is never repeated for both targets (as in our data), induces spurious spatial tuning for target 2 (slanted orange line) opposite to the true tuning for target 1. **C,D** Lasso regression coefficients across target locations for the same simulated neuron, under a balanced (**C**) and non-independent (**D**) design. Regression decorrelates responses to target 1 and 2, eliminating the spurious spatial tuning induced by the non-independent design (flat orange line in D). **E, F** Average firing rates (**E**) and firing rates estimated using Lasso regression (**F**) for three example neurons. Green lines represent T1 and orange lines represent T2. Regression-estimated coefficients from Lasso removed an anticorrelation bias in average firing rates likely due to the unbalanced task design.

**Supplementary Figure S4.**
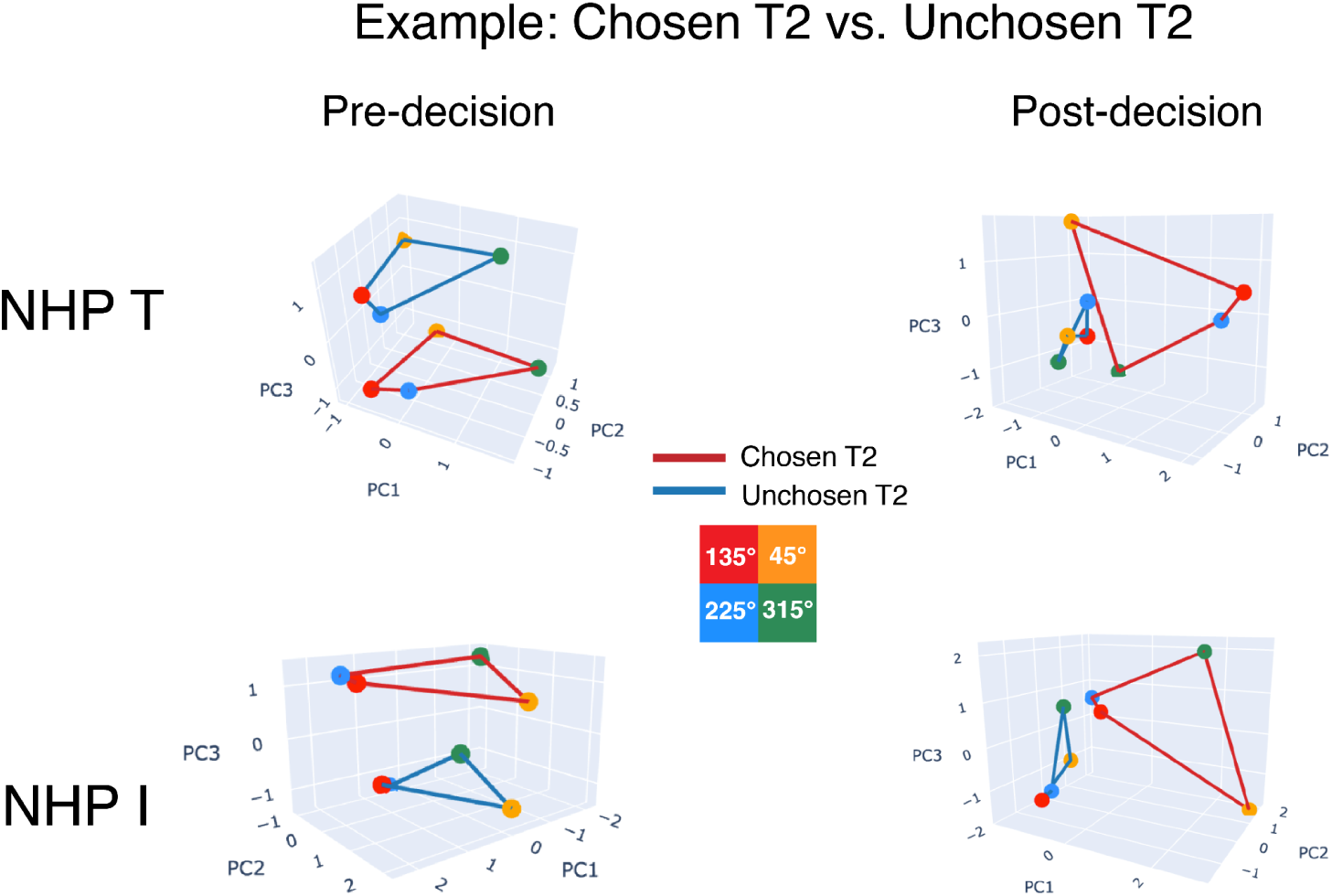
Representational geometry of *second* targets also showed decision-related rotation of chosen and unchosen targets into orthogonal subspaces. Analogous results to Figure 4C, but for the *second* (rather than the first) target. Geometry of chosen (red lines) and unchosen (blue lines) second targets before (left) and after (right) decision in reduced 3D space. Four colored dots correspond to the four target locations for each condition. After decision, chosen and unchosen targets are rotated into orthogonal subspaces.

**Supplementary Figure S5.**
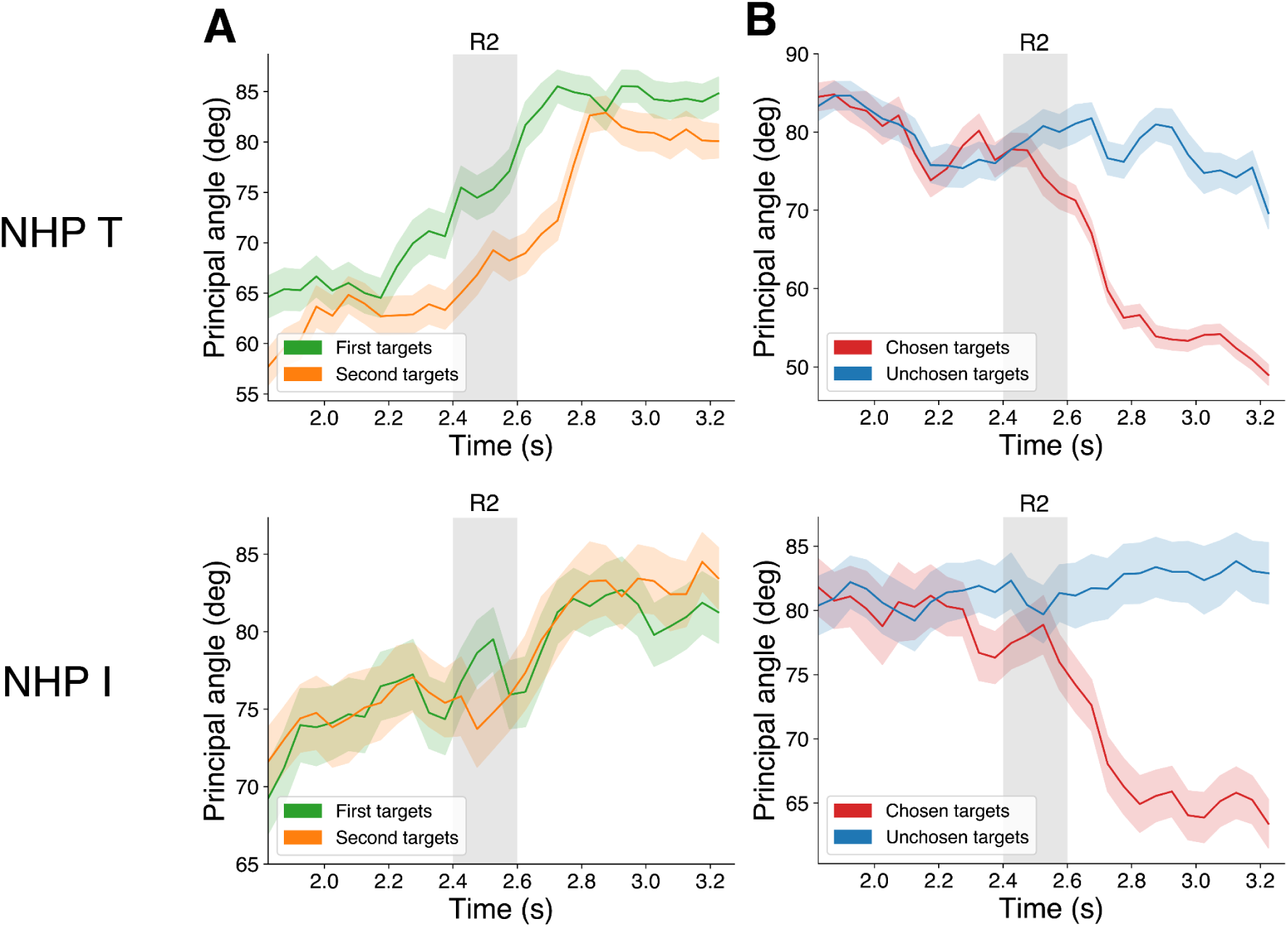
Principal angles showed similar results to subspace alignment. **A** The smallest principal angle between the first-target subspaces on trials when it was chosen vs unchosen (green line) and between the the second-target subspaces on trials when it was chosen vs unchosen (orange line) before and after decisions for both NHPs (cf. Figure 4D). **B** The smallest principal angle between the chosen-target subspaces on trials when the first vs second target was chosen (red line) and between the unchosen-target subspaces on trials when the first vs second target was unchosen (blue line) before and after decisions for both NHPs (cf. Figure 5D). Note that, in contrast to Figure 4D and 5D, here smaller values indicate stronger alignment, while 90 deg reflects perfect orthogonality.

**Supplementary Figure S6.**
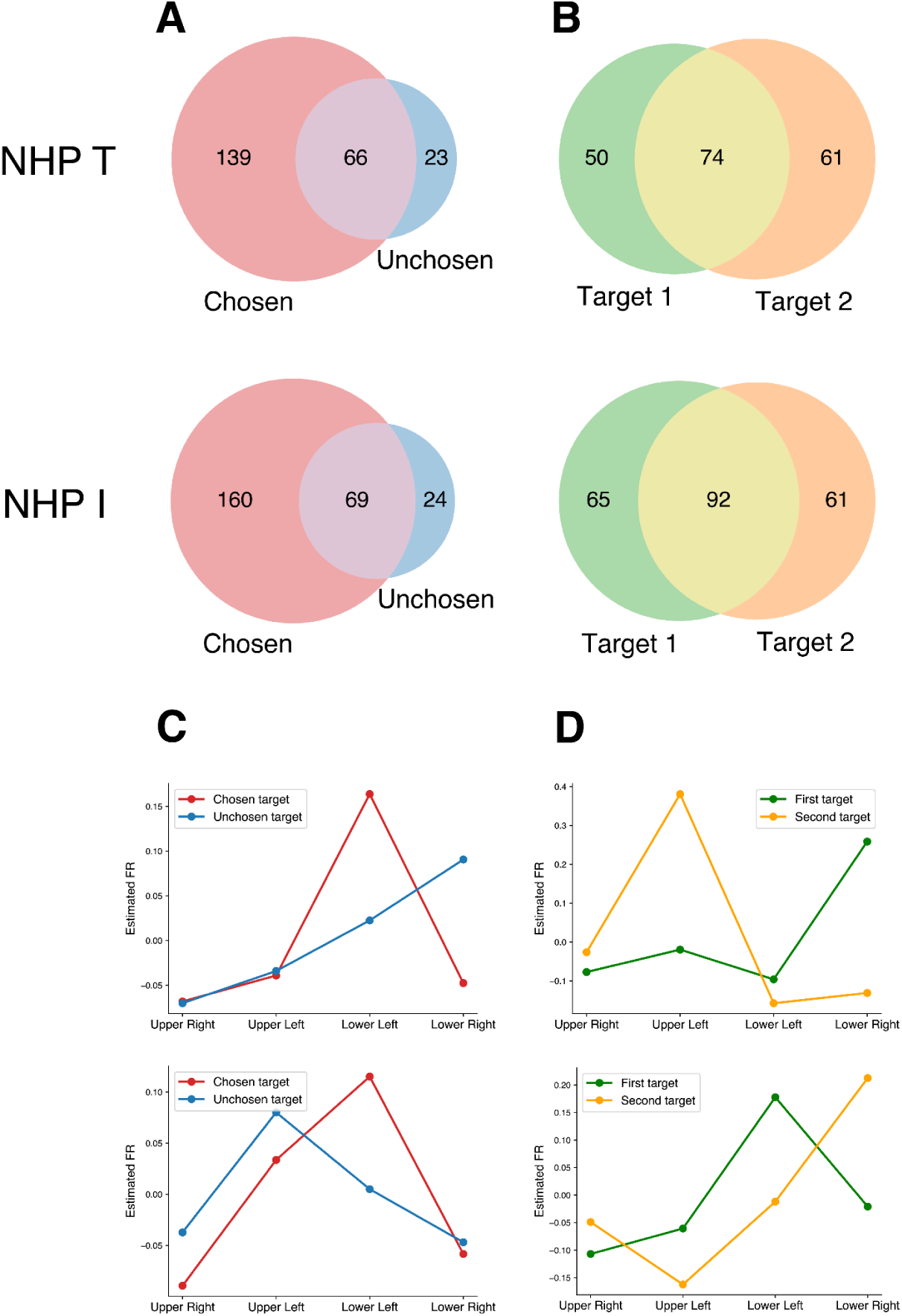
Orthogonal population geometry largely reflected overlapping neuronal populations rather than separate subpopulations. **A** Venn diagrams showing the number of neurons selective for the spatial location of the chosen target (red), unchosen target (blue), or both (purple), after the decision for each NHP (cf. post-decision orthogonal subspaces in Figure 4D). **B** Venn diagrams showing the number of neurons selective for the spatial location of the first target (green), second target (orange), or both (yellow), before the decision for each NHP (cf. pre-decision orthogonal subspaces in Figure 5D). **C, D** Estimated mean normalized firing rates of two example single neurons across the four target locations, for each choice status (chosen vs. unchosen targets) (**C**) and each sequential order (first vs. second targets) (**D**).

**Supplementary Figure S7.**
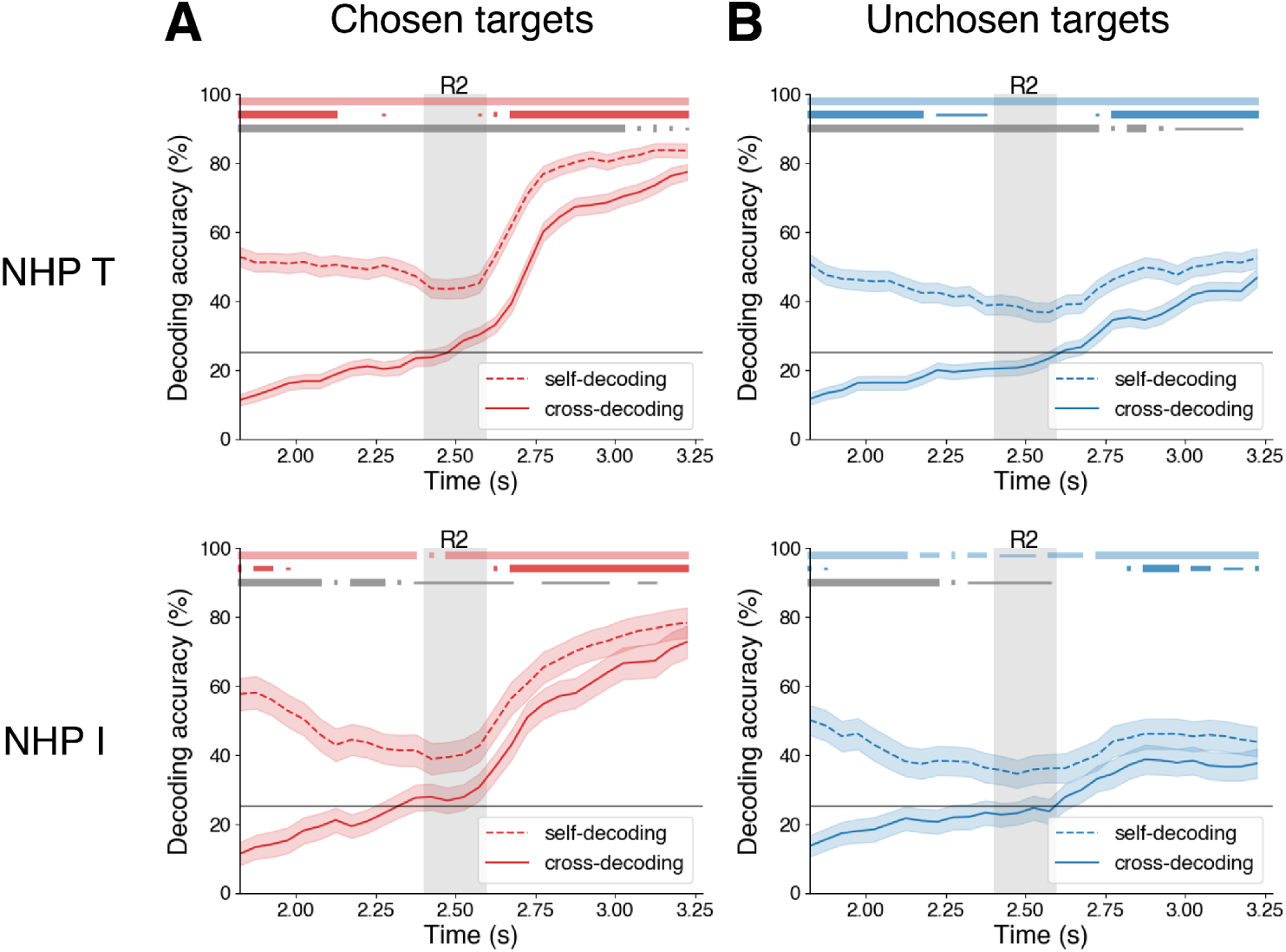
A single decoder can read out chosen target location after decisions, regardless of its initial presentation order. **A, B** Self- and cross-decoding accuracy for chosen targets (**A**), unchosen targets (**B**). Dashed lines are the self-decoding accuracies, where classifiers were tested on the same target order (first target/T1 vs second target/T2) that they were trained on (i.e. train T1→test T1, train T2→test T2). Solid lines are the cross-decoding accuracies when classifiers were tested on the opposite target order from the one they were trained on (i.e. train T1→test T2, train T2→test T1). Three series of markers from top to bottom represent significance of self-decoding accuracy against chance, cross-decoding accuracy against chance, and difference between self-decoding accuracy and cross-decoding accuracy (corrected bootstrap tests for all). Marker width indicates significance: p<0.05, p<0.01, p<0.001 for thin, medium, thick respectively. After the decision (R2), cross-decoding accuracy for chosen targets was well above chance, and similar to self-decoding accuracy. This suggests chosen target location could be read out by downstream areas using a single linear decoder, regardless of the chosen target’s initial presentation order. Similar results were found for unchosen targets, though the overall decoding accuracy is much lower.

**Supplementary Figure S8.**
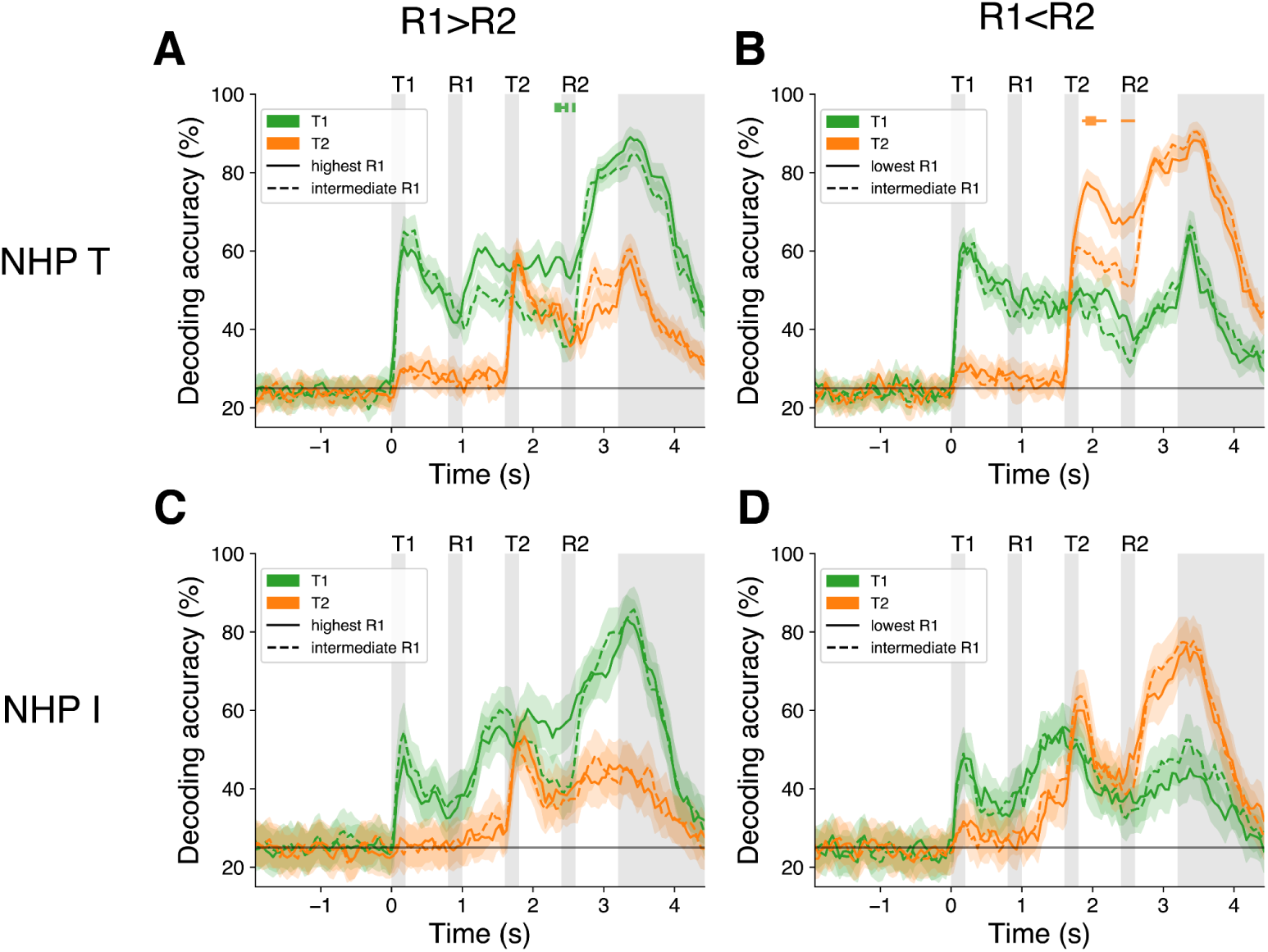
Evidence for early enhancement of location information for trials with highest/lowest first reward values. **A,C** For trials in which target 1 is chosen (R1>R2), there is elevated information reflecting the location of target 1 (T1) when reward value 1 (R1) is the highest possible value (solid green line) compared to when R1 is at an intermediate level (dashed green line). Markers at top represent the statistical significance difference between target 1 in two conditions (green) and target 2 in two conditions (orange). Marker width indicates significance: p<0.05, p<0.01, p<0.001 for thin, medium, thick respectively. There is a significant effect in NHP T that begins at the R1 presentation. (**A**) and a non-significant trend in NHP I (**C**). **B,D** For trials in which target 2 is chosen (R1<R2), for NHP T (**B**), there is elevated information about target 2 location (T2) when reward value 1 (R1) is the lowest compared to when R1 is at intermediate level. This begins as soon as information about the T2 location is available. For NHP I (**D**), there is no elevated T2 information. These results suggest that, at least for NHP T, when the highest or lowest possible reward value is offered first, the enhancement of location information reflecting the decision can happen immediately.

**Supplementary Figure S9.**
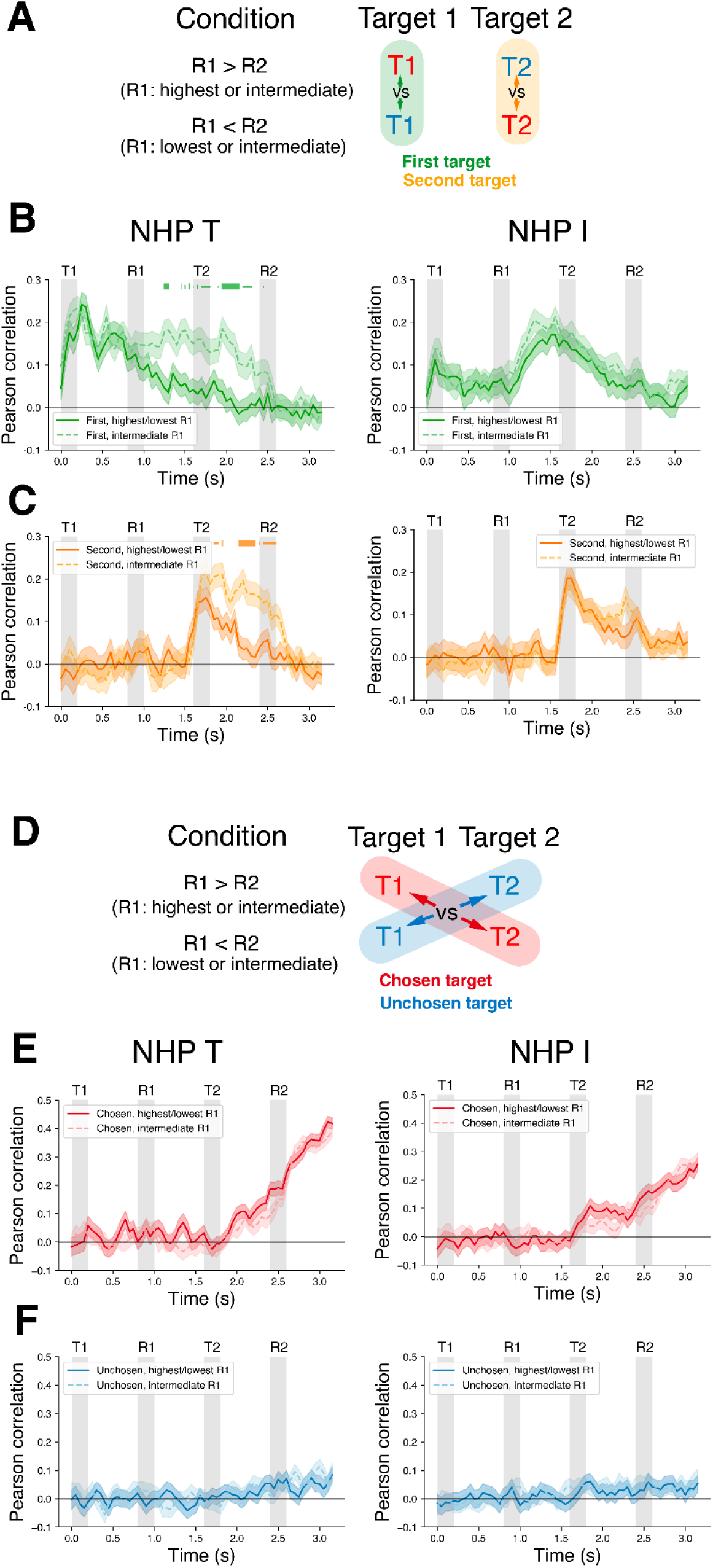
Evidence for early subspace reorganization for trials with the highest/lowest first reward trials. **A** Schematic showing the contrast between targets with the same order of presentation but resulting in different choices. We compared the representation of the first target on trials when it was chosen vs unchosen, separately for trials when the first reward cue (R1) signaled the highest/lowest possible values and for trials when R1 signaled the intermediate values. We also compared the representation of the second targets similarly. **B,C** Subspace alignment between the first target conditions (**B)** and between the second target conditions (**C**). Subspace alignment when R1 had intermediate values are plotted in light-colored lines, while subspace alignment when R1 had the highest/lowest values are plotted in dark-colored lines. Markers at top represent the statistical significance of the difference of subspace alignment between intermediate and highest/lowest trials. Marker width indicates significance: p<0.05, p<0.01, p<0.001 for thin, medium, thick respectively. There is a significant difference of subspace alignment for first and second targets for NHP T, but only non-significant trends for NHP I. **D** Schematic showing the contrast between targets with different order of presentation but resulting in the same choice (each chosen or each unchosen). We compared the representation of chosen targets when the first vs second target was chosen, separately for trials where the first reward cue (R1) signaled the highest/lowest possible values and for trials when R1 signaled the intermediate values. We also compared the representation of unchosen targets similarly. **E,F** Subspace alignment between the chosen targets (**E**), and between the unchosen targets (**F**). Both NHPs showed a small, non-significant difference of subspace alignment for chosen targets and no difference of subspace alignment for unchosen targets. These results provide evidence that, at least for NHP T, the subspace reorganization reflecting the decision can happen immediately when the highest or lowest possible reward value is offered first.

**Supplementary Figure S10.**
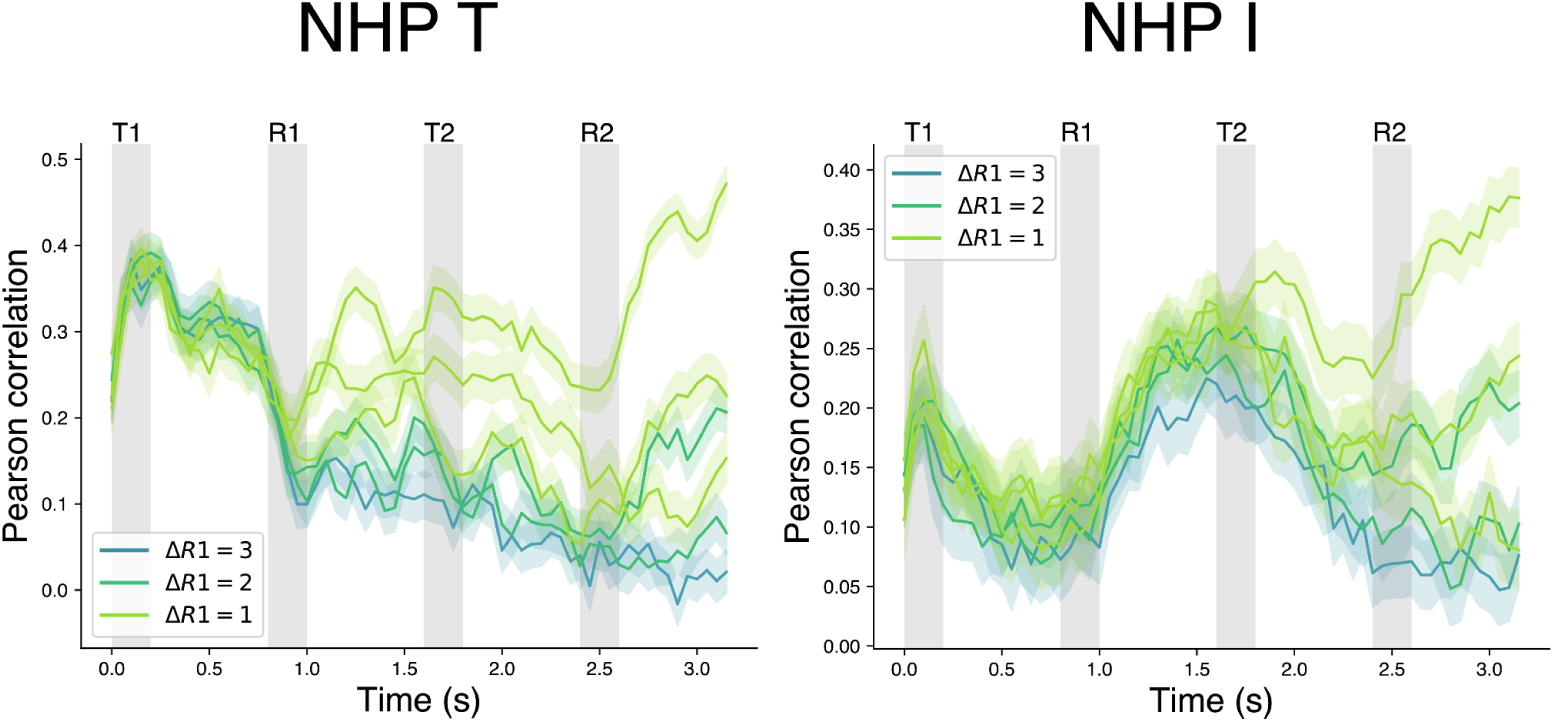
First targets increased alignment with more similar values. Alignment (correlation) between subspaces estimated for the first-presented spatial target (T1) for trials where the first reward value (R1) differed (ΔR1) by 1 (green), 2 (aqua), or 3 (blue) levels for both NHPs.

**Supplementary Figure S11.**
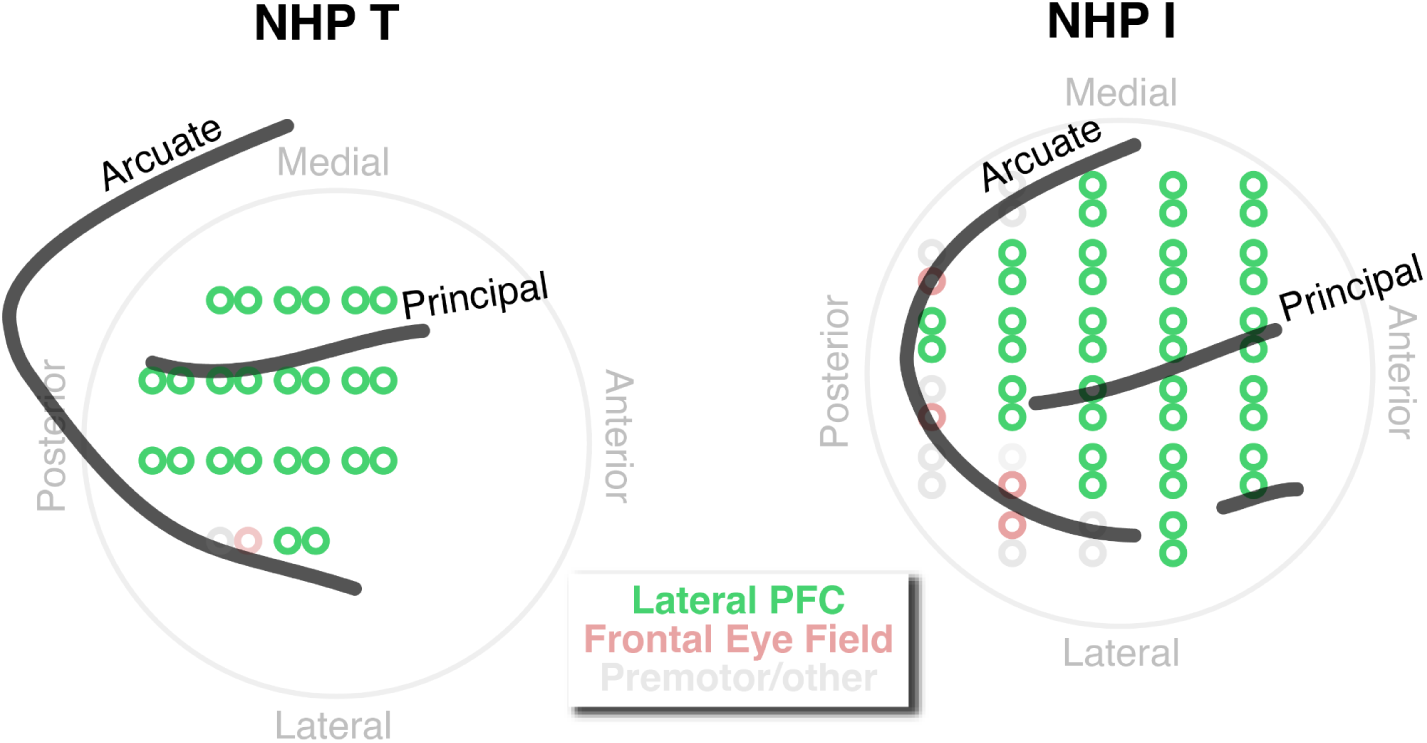
Lateral PFC recording sites. Maps of recording sites in each NHP subject across the cortical surface. Recording sites (green) spanned a wide region of right ventrolateral and dorsolateral PFC. Putative Frontal Eye Field (FEF) sites (red) were identified where low-current microstimulation evoked saccades. Excluding this small number of sites, as well as a few sites further posterior (gray), from analysis produced nearly identical results (Figure S12).

**Supplementary Figure S12.**
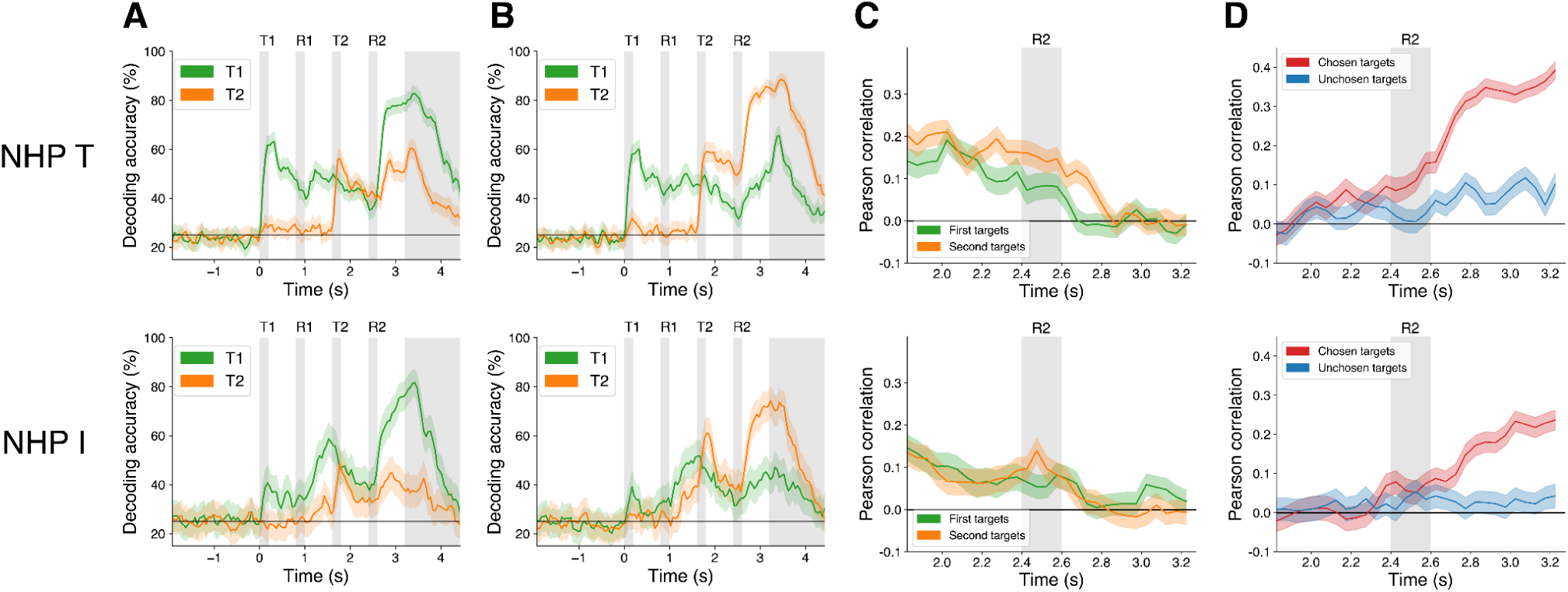
Results were nearly identical when reevaluated excluding putative Frontal Eye Field (FEF)/premotor cortex (PMC) sites. **A, B** Decoding accuracy of first (green) and second (orange) targets when R1>R2 (**A**) and when R1<R2 (**B**) when the small number of putative FEF/PMC sites were removed (compare to Figure 2B,C). **C, D** Subspace alignment between targets of the same order of presentation (**C**) and between targets of the same choice (**D**) when FEF/PMC sites were removed (compare to Figure 4D,5D).

## References

1. Kable JW, Glimcher PW. The Neurobiology of Decision: Consensus and Controversy. Neuron. 2009 Sep 24;63(6):733–45. doi:10.1016/j.neuron.2009.09.003

2. Padoa-Schioppa C. Neurobiology of Economic Choice: A Good-Based Model. Annu Rev Neurosci. 2011 Jul 21;34(Volume 34, 2011):333–59. doi:10.1146/annurev-neuro-061010-113648

3. Rangel A, Camerer C, Montague PR. A framework for studying the neurobiology of value-based decision making. Nat Rev Neurosci. 2008 Jul;9(7):545–56. doi:10.1038/nrn2357

4. Rangel A, Hare T. Neural computations associated with goal-directed choice. Curr Opin Neurobiol. 2010 Apr 1;Cognitive neuroscience20(2):262–70. doi:10.1016/j.conb.2010.03.001

5. Yoo SBM, Hayden BY. Economic Choice as an Untangling of Options into Actions. Neuron. 2018 Aug 8;99(3):434–47. doi:10.1016/j.neuron.2018.06.038

6. Ballesta S, Shi W, Conen KE, Padoa-Schioppa C. Values encoded in orbitofrontal cortex are causally related to economic choices. Nature. 2020 Dec;588(7838):450–3. doi:10.1038/s41586-020-2880-x

7. Ballesta S, Shi W, Padoa-Schioppa C. Orbitofrontal cortex contributes to the comparison of values underlying economic choices. Nat Commun. 2022 Jul 29;13(1):4405. doi:10.1038/s41467-022-32199-y

8. Kimmel DL, Elsayed GF, Cunningham JP, Newsome WT. Value and choice as separable and stable representations in orbitofrontal cortex. Nat Commun. 2020 Jul 10;11(1):1. doi:10.1038/s41467-020-17058-y

9. Strait CE, Blanchard TC, Hayden BY. Reward Value Comparison via Mutual Inhibition in Ventromedial Prefrontal Cortex. Neuron. 2014 Jun 18;82(6):1357–66. doi:10.1016/j.neuron.2014.04.032 PubMed PMID: 24881835.

10. Seo M, Lee E, Averbeck BB. Action Selection and Action Value in Frontal-Striatal Circuits. Neuron. 2012 Jun 7;74(5):947–60. doi:10.1016/j.neuron.2012.03.037 PubMed PMID: 22681697.

11. Cai X, Padoa-Schioppa C. Contributions of Orbitofrontal and Lateral Prefrontal Cortices to Economic Choice and the Good-to-Action Transformation. Neuron. 2014 Mar 5;81(5):1140–51. doi:10.1016/j.neuron.2014.01.008

12. Yim MY, Cai X, Wang XJ. Transforming the Choice Outcome to an Action Plan in Monkey Lateral Prefrontal Cortex: A Neural Circuit Model. Neuron. 2019 Aug 7;103(3):520–532.e5. doi:10.1016/j.neuron.2019.05.032 PubMed PMID: 31230761.

13. Kennerley SW, Wallis JD. Reward-Dependent Modulation of Working Memory in Lateral Prefrontal Cortex. J Neurosci. 2009 Mar 11;29(10):3259–70. doi:10.1523/JNEUROSCI.5353-08.2009 PubMed PMID: 19279263.

14. Petrides M, Pandya DN. Efferent association pathways originating in the caudal prefrontal cortex in the macaque monkey. J Comp Neurol. 2006 Sep 10;498(2):227–51. doi:10.1002/cne.21048 PubMed PMID: 16856142.

15. Saleem KS, Miller B, Price JL. Subdivisions and connectional networks of the lateral prefrontal cortex in the macaque monkey. J Comp Neurol. 2014 May;522(7):1641–90. doi:10.1002/cne.23498

16. Wallis JD. Orbitofrontal Cortex and Its Contribution to Decision-Making. Annu Rev Neurosci. 2007 Jul 21;30(Volume 30, 2007):31–56. doi:10.1146/annurev.neuro.30.051606.094334

17. Lu MT, Preston JB, Strick PL. Interconnections between the prefrontal cortex and the premotor areas in the frontal lobe. J Comp Neurol. 1994;341(3):375–92. doi:10.1002/cne.903410308

18. Takada M, Nambu A, Hatanaka N, Tachibana Y, Miyachi S, Taira M, et al. Organization of prefrontal outflow toward frontal motor-related areas in macaque monkeys. Eur J Neurosci. 2004;19(12):3328–42. doi:10.1111/j.0953-816X.2004.03425.x

19. Takahara D, Inoue K ichi, Hirata Y, Miyachi S, Nambu A, Takada M, et al. Multisynaptic projections from the ventrolateral prefrontal cortex to the dorsal premotor cortex in macaques – anatomical substrate for conditional visuomotor behavior. Eur J Neurosci. 2012;36(10):3365–75. doi:10.1111/j.1460-9568.2012.08251.x

20. Mante V, Sussillo D, Shenoy KV, Newsome WT. Context-dependent computation by recurrent dynamics in prefrontal cortex. Nature. 2013 Nov;503(7474):78–84. doi:10.1038/nature12742

21. Abbass M, Corrigan B, Johnston R, Gulli R, Sachs A, Lau JC, et al. Prefrontal cortex neuronal ensembles dynamically encode task features during associative memory and virtual navigation. Cell Rep. 2025 Jan 28;44(1). doi:10.1016/j.celrep.2024.115124 PubMed PMID: 39772389.

22. Weber J, Iwama G, Solbakk AK, Blenkmann AO, Larsson PG, Ivanovic J, et al. Subspace partitioning in the human prefrontal cortex resolves cognitive interference. Proc Natl Acad Sci. 2023 Jul 11;120(28):e2220523120. doi:10.1073/pnas.2220523120

23. Xie Y, Hu P, Li J, Chen J, Song W, Wang XJ, et al. Geometry of sequence working memory in macaque prefrontal cortex. Science. 2022 Feb 11;375(6581):632–9. doi:10.1126/science.abm0204

24. Tian Z, Chen J, Zhang C, Min B, Xu B, Wang L. Mental programming of spatial sequences in working memory in the macaque frontal cortex. Science. 2024 Sep 27;385(6716):eadp6091. doi:10.1126/science.adp6091

25. Tang C, Herikstad R, Parthasarathy A, Libedinsky C, Yen SC. Minimally dependent activity subspaces for working memory and motor preparation in the lateral prefrontal cortex. Serences JT, Frank MJ, editors. eLife. 2020 Sep 9;9:e58154. doi:10.7554/eLife.58154

26. Parthasarathy A, Tang C, Herikstad R, Cheong LF, Yen SC, Libedinsky C. Time-invariant working memory representations in the presence of code-morphing in the lateral prefrontal cortex. Nat Commun. 2019 Nov 1;10(1):4995. doi:10.1038/s41467-019-12841-y

27. Rouzitalab A, Boulay CB, Park J, Martinez-Trujillo JC, Sachs AJ. Ensembles code for associative learning in the primate lateral prefrontal cortex. Cell Rep. 2023 May 30;42(5). doi:10.1016/j.celrep.2023.112449 PubMed PMID: 37119136.

28. Aoi MC, Mante V, Pillow JW. Prefrontal cortex exhibits multidimensional dynamic encoding during decision-making. Nat Neurosci. 2020 Nov;23(11):1410–20. doi:10.1038/s41593-020-0696-5

29. Murray JD, Bernacchia A, Roy NA, Constantinidis C, Romo R, Wang XJ. Stable population coding for working memory coexists with heterogeneous neural dynamics in prefrontal cortex. Proc Natl Acad Sci. 2017 Jan 10;114(2):394–9. doi:10.1073/pnas.1619449114

30. Churchland MM, Cunningham JP, Kaufman MT, Foster JD, Nuyujukian P, Ryu SI, et al. Neural population dynamics during reaching. Nature. 2012 Jul;487(7405):51–6. doi:10.1038/nature11129

31. Elsayed GF, Lara AH, Kaufman MT, Churchland MM, Cunningham JP. Reorganization between preparatory and movement population responses in motor cortex. Nat Commun. 2016 Oct 27;7(1):1. doi:10.1038/ncomms13239

32. Perich MG, Gallego JA, Miller LE. A Neural Population Mechanism for Rapid Learning. Neuron. 2018 Nov;100(4):964–976.e7. doi:10.1016/j.neuron.2018.09.030

33. Sadtler PT, Quick KM, Golub MD, Chase SM, Ryu SI, Tyler-Kabara EC, et al. Neural constraints on learning. Nature. 2014 Aug;512(7515):423–6. doi:10.1038/nature13665

34. Ruff DA, Ni AM, Cohen MR. Cognition as a Window into Neuronal Population Space. Annu Rev Neurosci. 2018 Jul 8;41(1):77–97. doi:10.1146/annurev-neuro-080317-061936

35. Ebitz RB, Hayden BY. The population doctrine in cognitive neuroscience. Neuron. 2021 Oct;109(19):3055–68. doi:10.1016/j.neuron.2021.07.011

36. Gallego JA, Perich MG, Miller LE, Solla SA. Neural Manifolds for the Control of Movement. Neuron. 2017 Jun;94(5):978–84. doi:10.1016/j.neuron.2017.05.025

37. Libby A, Buschman TJ. Rotational dynamics reduce interference between sensory and memory representations. Nat Neurosci. 2021 May;24(5):5. doi:10.1038/s41593-021-00821-9

38. Panichello MF, Buschman TJ. Shared mechanisms underlie the control of working memory and attention. Nature. 2021 Apr;592(7855):7855. doi:10.1038/s41586-021-03390-w

39. Flesch T, Juechems K, Dumbalska T, Saxe A, Summerfield C. Orthogonal representations for robust context-dependent task performance in brains and neural networks. Neuron. 2022 Apr 6;110(7):1258–1270.e11. doi:10.1016/j.neuron.2022.01.005 PubMed PMID: 35085492.

40. Kaufman MT, Churchland MM, Ryu SI, Shenoy KV. Cortical activity in the null space: permitting preparation without movement. Nat Neurosci. 2014 Mar;17(3):3. doi:10.1038/nn.3643

41. Semedo JD, Jasper AI, Zandvakili A, Krishna A, Aschner A, Machens CK, et al. Feedforward and feedback interactions between visual cortical areas use different population activity patterns. Nat Commun. 2022 Mar 1;13(1):1099. doi:10.1038/s41467-022-28552-w

42. MacDowell CJ, Libby A, Jahn CI, Tafazoli S, Ardalan A, Buschman TJ. Multiplexed subspaces route neural activity across brain-wide networks. Nat Commun. 2025 Apr 9;16(1):3359. doi:10.1038/s41467-025-58698-2

43. Stokes MG, Kusunoki M, Sigala N, Nili H, Gaffan D, Duncan J. Dynamic coding for cognitive control in prefrontal cortex. Neuron. 2013 Apr 24;78(2):364–75. doi:10.1016/j.neuron.2013.01.039 PubMed PMID: 23562541; PubMed Central PMCID: PMC3898895.

44. Bernardi S, Benna MK, Rigotti M, Munuera J, Fusi S, Salzman CD. The Geometry of Abstraction in the Hippocampus and Prefrontal Cortex. Cell. 2020 Nov 12;183(4):954–967.e21. doi:10.1016/j.cell.2020.09.031

45. Gallego JA, Perich MG, Naufel SN, Ethier C, Solla SA, Miller LE. Cortical population activity within a preserved neural manifold underlies multiple motor behaviors. Nat Commun. 2018 Oct 12;9(1):4233. doi:10.1038/s41467-018-06560-z

46. Bouchacourt F, Buschman TJ. A Flexible Model of Working Memory. Neuron. 2019 Jul;103(1):147–160.e8. doi:10.1016/j.neuron.2019.04.020

47. Nogueira R, Rodgers CC, Bruno RM, Fusi S. The geometry of cortical representations of touch in rodents. Nat Neurosci. 2023 Feb;26(2):239–50. doi:10.1038/s41593-022-01237-9

48. Ritz H, Shenhav A. Orthogonal neural encoding of targets and distractors supports multivariate cognitive control. Nat Hum Behav. 2024 Mar 8;1–17. doi:10.1038/s41562-024-01826-7

49. Srinath R, Ni AM, Marucci C, Cohen MR, Brainard DH. Orthogonal neural representations support perceptual judgements of natural stimuli [Internet]. bioRxiv; 2024 [cited 2024 Apr 26]. p. 2024.02.14.580134. Available from: https://www.biorxiv.org/content/10.1101/2024.02.14.580134v1 doi:10.1101/2024.02.14.580134

50. Charlton JA, Goris RLT. Abstract deliberation by visuomotor neurons in prefrontal cortex. Nat Neurosci. 2024 Jun;27(6):1167–75. doi:10.1038/s41593-024-01635-1

51. Johnston WJ, Fine JM, Yoo SBM, Ebitz RB, Hayden BY. Semi-orthogonal subspaces for value mediate a binding and generalization trade-off. Nat Neurosci. 2024 Sep 17;1–13. doi:10.1038/s41593-024-01758-5

52. Ito T, Klinger T, Schultz DH, Murray JD, Cole MW, Rigotti M. Compositional generalization through abstract representations in human and artificial neural networks [Internet]. arXiv; 2022 [cited 2026 May 21]. Available from: https://arxiv.org/abs/2209.07431 doi:10.48550/ARXIV.2209.07431

53. Luyckx F, Nili H, Spitzer B, Summerfield C. Neural structure mapping in human probabilistic reward learning. Lee D, Gold JI, Lee D, Chafee M, editors. eLife. 2019 Mar 7;8:e42816. doi:10.7554/eLife.42816

54. Morton NW, Schlichting ML, Preston AR. Representations of common event structure in medial temporal lobe and frontoparietal cortex support efficient inference. Proc Natl Acad Sci. 2020 Nov 24;117(47):29338–45. doi:10.1073/pnas.1912338117

55. Muhle-Karbe PS, Sheahan H, Pezzulo G, Spiers HJ, Chien S, Schuck NW, et al. Goal-seeking compresses neural codes for space in the human hippocampus and orbitofrontal cortex. Neuron. 2023 Sep;S0896627323006323. doi:10.1016/j.neuron.2023.08.021

56. Sheahan H, Luyckx F, Nelli S, Teupe C, Summerfield C. Neural state space alignment for magnitude generalization in humans and recurrent networks. Neuron. 2021 Apr 7;109(7):1214–1226.e8. doi:10.1016/j.neuron.2021.02.004 PubMed PMID: 33626322.

57. Asaad WF, Lauro PM, Perge JA, Eskandar EN. Prefrontal Neurons Encode a Solution to the Credit-Assignment Problem. J Neurosci. 2017 Jul 19;37(29):6995–7007. doi:10.1523/JNEUROSCI.3311-16.2017 PubMed PMID: 28634307.

58. Yoo SBM, Hayden BY. The Transition from Evaluation to Selection Involves Neural Subspace Reorganization in Core Reward Regions. Neuron. 2020 Feb 19;105(4):712–724.e4. doi:10.1016/j.neuron.2019.11.013

59. Elsayed GF, Lara AH, Kaufman MT, Churchland MM, Cunningham JP. Reorganization between preparatory and movement population responses in motor cortex. Nat Commun. 2016 Oct;27;7(1):13239.

60. Rigotti M, Barak O, Warden MR, Wang XJ, Daw ND, Miller EK, et al. The importance of mixed selectivity in complex cognitive tasks. Nature. 2013 May;497(7451):7451. doi:10.1038/nature12160

61. Buschman TJ, Kastner S. From Behavior to Neural Dynamics: An Integrated Theory of Attention. Neuron. 2015 Oct 7;88(1):127–44. doi:10.1016/j.neuron.2015.09.017 PubMed PMID: 26447577.

62. Desimone R, Duncan J. Neural Mechanisms of Selective Visual Attention. Annu Rev Neurosci. 1995 Mar 1;18(Volume 18, 1995):193–222. doi:10.1146/annurev.ne.18.030195.001205

63. Hasegawa RP, Matsumoto M, Mikami A. Search Target Selection in Monkey Prefrontal Cortex. J Neurophysiol. 2000 Sep;84(3):1692–6. doi:10.1152/jn.2000.84.3.1692

64. McAdams CJ, Reid RC. Attention Modulates the Responses of Simple Cells in Monkey Primary Visual Cortex. J Neurosci. 2005 Nov 23;25(47):11023–33. doi:10.1523/JNEUROSCI.2904-05.2005 PubMed PMID: 16306415.

65. Reynolds JH, Pasternak T, Desimone R. Attention Increases Sensitivity of V4 Neurons. Neuron. 2000 Jun 1;26(3):703–14. doi:10.1016/S0896-6273(00)81206-4 PubMed PMID: 10896165.

66. Reynolds JH, Heeger DJ. The Normalization Model of Attention. Neuron. 2009 Jan 29;61(2):168–85. doi:10.1016/j.neuron.2009.01.002 PubMed PMID: 19186161.

67. Suzuki M, Gottlieb J. Distinct neural mechanisms of distractor suppression in the frontal and parietal lobe. Nat Neurosci. 2013;Jan;16(1):98–104.

68. Cheng YA, Sanayei M, Chen X, Jia K, Li S, Fang F, et al. A neural geometry approach comprehensively explains apparently conflicting models of visual perceptual learning. Nat Hum Behav. 2025 Mar 31;9(5):1023–40. doi:10.1038/s41562-025-02149-x

69. Saxena S, Cunningham JP. Towards the neural population doctrine. Curr Opin Neurobiol. 2019 Apr;55:103–11. doi:10.1016/j.conb.2019.02.002

70. Lee D, Seo H, Jung MW. Neural Basis of Reinforcement Learning and Decision Making. Annu Rev Neurosci. 2012 Jul 21;35(Volume 35, 2012):287–308. doi:10.1146/annurev-neuro-062111-150512

71. Kennerley SW, Dahmubed AF, Lara AH, Wallis JD. Neurons in the Frontal Lobe Encode the Value of Multiple Decision Variables. J Cogn Neurosci. 2009 Jun 1;21(6):1162–78. doi:10.1162/jocn.2009.21100

72. Lim SL, O’Doherty JP, Rangel A. The Decision Value Computations in the vmPFC and Striatum Use a Relative Value Code That is Guided by Visual Attention. J Neurosci. 2011 Sep 14;31(37):13214–23. doi:10.1523/JNEUROSCI.1246-11.2011 PubMed PMID: 21917804.

73. Padoa-Schioppa C, Assad JA. Neurons in the orbitofrontal cortex encode economic value. Nature. 2006 May;441(7090):223–6. doi:10.1038/nature04676

74. Roesch MR, Olson CR. Neuronal Activity in Primate Orbitofrontal Cortex Reflects the Value of Time. J Neurophysiol. 2005 Oct;94(4):2457–71. doi:10.1152/jn.00373.2005

75. Tremblay L, Schultz W. Relative reward preference in primate orbitofrontal cortex. Nature. 1999 Apr;398(6729):704–8. doi:10.1038/19525

76. Wunderlich K, Rangel A, O’Doherty JP. Economic choices can be made using only stimulus values. Proc Natl Acad Sci. 2010 Aug 24;107(34):15005–10. doi:10.1073/pnas.1002258107

77. Munet NT, Wallis JD. Effects of overt and covert attention on decision-making dynamics in prefrontal cortex [Internet]. bioRxiv; 2026 [cited 2026 Jul 1]. p. 2026.05.18.723036. Available from: https://www.biorxiv.org/content/10.64898/2026.05.18.723036v1 doi:10.64898/2026.05.18.723036

78. Chen X, Stuphorn V. Sequential selection of economic good and action in medial frontal cortex of macaques during value-based decisions. Schultz W, editor. eLife. 2015 Nov 27;4:e09418. doi:10.7554/eLife.09418

79. Sul JH, Jo S, Lee D, Jung MW. Role of rodent secondary motor cortex in value-based action selection. Nat Neurosci. 2011 Aug 14;14(9):1202–8. doi:10.1038/nn.2881 PubMed PMID: 21841777; PubMed Central PMCID: PMC3164897.

80. Balewski ZZ, Elston TW, Knudsen EB, Wallis JD. Value dynamics affect choice preparation during decision-making. Nat Neurosci. 2023 Aug 10;1–9. doi:10.1038/s41593-023-01407-3

81. Kobayashi S, Nomoto K, Watanabe M, Hikosaka O, Schultz W, Sakagami M. Influences of Rewarding and Aversive Outcomes on Activity in Macaque Lateral Prefrontal Cortex. Neuron. 2006 Sep 21;51(6):861–70. doi:10.1016/j.neuron.2006.08.031 PubMed PMID: 16982429.

82. Wallis JD, Miller EK. Neuronal activity in primate dorsolateral and orbital prefrontal cortex during performance of a reward preference task. Eur J Neurosci. 2003;18(7):2069–81. doi:10.1046/j.1460-9568.2003.02922.x

83. Meyers EM, Kreiman G. Tutorial on pattern classification in cell recording. In: Visual Population Codes. MIT Press; 2012. p. 517–38.

84. Efron B, Tibshirani RJ. An Introduction to the Bootstrap [Internet]. 0 ed. Chapman and Hall/CRC; 1994 [cited 2026 Jan 28]. Available from: https://www.taylorfrancis.com/books/9781000064988 doi:10.1201/9780429246593

85. Dobs K, Yuan J, Martinez J, Kanwisher N. Behavioral signatures of face perception emerge in deep neural networks optimized for face recognition. Proc Natl Acad Sci. 2023 Aug 8;120(32):e2220642120. doi:10.1073/pnas.2220642120

86. Fukuda H, Ma N, Suzuki S, Harasawa N, Ueno K, Gardner JL, et al. Computing Social Value Conversion in the Human Brain. J Neurosci. 2019 Jun 26;39(26):5153–72. doi:10.1523/JNEUROSCI.3117-18.2019

87. Benjamini Y, Hochberg Y. Controlling the false discovery rate: a practical and powerful approach to multiple testing. J R Stat Soc Ser B Methodol. 1995;289–300.

